# Cytoplasmic Colocalization of Granulins and TDP-43 Prion-like Domain Involves Electrostatically Driven Coacervation Tuned by the Redox State of Cysteines

**DOI:** 10.1101/2021.06.25.449959

**Authors:** Anukool A. Bhopatkar, Shailendra Dhakal, Vijayaraghavan Rangachari

**Affiliations:** Department of Chemistry and Biochemistry, School of Mathematics and Natural Sciences, University of Southern Mississippi, Hattiesburg, MS 39406; Center for Molecular and Cellular Biosciences, University of Southern Mississippi, Hattiesburg, MS 39406

**Keywords:** granulin, progranulin, TDP-43, FTLD, ALS, liquid-liquid phase separation, stress granules

## Abstract

Cytoplasmic inclusions containing aberrant proteolytic fragments of TDP-43 are associated with frontotemporal lobar degeneration (FTLD) and other related pathologies. In FTLD, TDP-43 is translocated into the cytoplasm and proteolytically cleaved to generate a prion-like domain (PrLD) containing C-terminal fragments (C25 and C35) that form toxic inclusions. Under stress, TDP-43 partitions into membraneless organelles called stress granules (SGs) by coacervating with RNA and other proteins. We were interested in understanding if and how cysteine-rich granulins (GRNs 1-7), which are the proteolytic products of a genetic risk factor in FTLD called progranulin, interact with TDP-43. We show that extracellular GRNs internalize and colocalize with PrLD as puncta in the cytoplasm of neuroblastoma cells but show no presence in SGs. In addition, we show GRNs and PrLD coacervate to undergo liquid-liquid phase separation (LLPS) or form gel- or solid-like aggregates. Identification of the sequence determinants within GRNs for the observed phase transitions reveal the negative charges to be the drivers of LLPS modulated by the positive charges and the redox state of cysteines. Furthermore, RNA and GRNs compete and expunge one another from PrLD condensates, providing a basis for GRN’s absence in SGs. Together, the results illustrate the potential mechanisms by which extracellular GRNs, formed during chronic inflammatory conditions, could internalize, and modulate cytoplasmic TDP-43 inclusions in proteinopathies.

## Introduction

Frontotemporal lobar degeneration (FTLD), is a progressive neurodegenerative disease that affects the frontal and temporal lobes of patients predominantly 45-65 years of age ^1^. The neuronal atrophy observed in these patients is associated with language impairments accompanied by behavioral and personality changes ^2–3^. Based on histopathological signatures, FTLD is subclassified into FTLD-TDP, FTLD-Tau, and FTLD-FUS (rare) ^1, 4–5^. Nearly half of FTLD patients show the presence of neuronal and glial cytoplasmic inclusions of TAR DNA binding protein-43 (TDP-43) ^5^, making FTLD-TDP a predominant category of the disease. Furthermore, 95% of sporadic amyotrophic lateral sclerosis (ALS) patients also show TDP-43 inclusions besides sharing common genetic etiologies with FTLD, making FTLD-TDP and ALS to be part of a clinicopathological continuum ^5^. Molecular etiologies of familial FTLD are associated with mutations in microtubule-associated protein tau (*MAPT*) for FTLD-Tau, and those in Progranulin (*GRN*), TDP-43 (*TARDBP*), Fused in sarcoma (*FUS*), and Hexanucleotide repeat expansion in C9orf72 (*C9orf72*), for FTLD-TDP ^2, 4, 6^.

TDP-43 is a 43 kDa protein that contains an N-terminal domain, two RNA recognition motifs (RRMs), and a prion-like disordered, C-terminal domain enriched with low complexity sequences (PrLD) ^4, 7^. Localized in the nucleus, TDP-43 is involved in a wide range of functions including transcriptional regulation, RNA metabolism and splicing among others ^8–10^. Under cellular stress such as heat shock, oxidative insults, inflammation ^11–12^ or nutrient starvation ^12^, TDP-43 coacervates with mRNA and other proteins to undergo liquid-liquid phase separation (LLPS) and partitions into membraneless organelles called stress granules (SGs) ^11, 13^. SGs are reversible cytoplasmic assemblies formed in response to cellular stress to sequester and prevent mRNA transcripts from degradation or translation ^14–16^. In pathological conditions, TDP-43 is translocated to the cytoplasm where it undergoes aberrant proteolytic cleavage to generate several C-terminal fragments (CTFs) of varying sizes including C35 (~35 kDa), C25 (~25 kDa), and C17 (~17 kDa) with the former two being the most abundant of them ^17–20^. These fragments are also known to be formed by alternative splicing to varying degrees ^21^. Importantly, TDP-43 CTFs undergo aggregation to form toxic, insoluble inclusions in the cytoplasm that are the pathological hallmarks of ALS and FTLD patients ^19, 22^. Despite the conspicuity of the TDP-43 CTF aggregates in patients, factors responsible for their biogenesis and subsequent roles in pathology remain poorly understood. Similarly, while the formation and functions of biomolecular condensates is well established for TDP-43 and other members of SGs such as hnRNP1 ^23–24^, FUS ^23, 25^ and TIA1 ^23, 26^, the link between SGs and amyloid formation remains ambiguous, but one that could be key in neurodegenerative pathologies.

Another key protein implicated in FTLD is a 68.5 kDa secreted protein called progranulin (PGRN). The protein possesses pleiotropic functions including wound healing, tumorigenesis ^27^ and immunomodulation ^28^. In neurons, PGRN has been identified to play key roles in neuronal functions, lysosomal homeostasis, cell survival, and differentiation ^29–35^, and has gained significant attention due to its link to neurodegenerative pathologies especially FTLD. About 30-50% of FTLD-TDP cases are of a heritable type with mutations in *GRN* underlying a majority of them ^36–37^. The autosomal dominant heterozygous *GRN* mutations results in haploinsufficiency of the protein, which led to the conclusion that PGRN plays a neuroprotective role ^38–40^, while homozygous *GRN* mutations lead to neuronal ceroid lipofuscinosis (NCL), a lysosomal disease ^39, 41^. But for the genetic connection and haploinsufficiency, the precise mechanism by which PGRN may influence FTLD remains unknown. In this context, the cysteine-rich, ~6 kDa modules called granulins (GRNs 1-7), which are the proteolytic products of PGRN ^42^ have been of great interest to us for their potential roles in FTLD and other related pathologies. It is now known that extracellular PGRN is endocytosed into the lysosome via a sortillin-mediated pathway to be processed by cathepsins into GRNs ^43^, which are thus speculated to possess lysosomal functions ^44–45^. GRNs also function in a plethora of roles in normal cell biology ^46–48^ but possess opposing inflammatory properties to PGRN; while PGRN is anti-inflammatory, GRNs show pro-inflammatory properties ^42^. During inflammation, PGRN secreted from activated microglia and astrocytes undergoes extracellular proteolysis by neutrophil elastases and other proteases to generate GRNs ^49^. The fate of the extracellular GRNs on neuronal function and dysfunction continues to remain undetermined and we conjecture that they are taken up by neurons where they interact with modulate TDP-43 assemblies. Some support for this idea came from a report from Salazar and coworkers who showed that specific GRNs affected TDP-43 toxicity and behavior functions in a *C.elegens* model ^50^. We also demonstrated that GRNs, 3 and 5, interact with and modulate the aggregation and phase behavior of TDP-43 PrLD *in vitro* ^51^. Furthermore, GRN immunopositivity observed in several regions of the brain in post-mortem Alzheimer disease and FTLD-TDP patients potentiates the significance of GRNs in pathology ^52^. Importantly, acute inflammation such as traumatic brain injury (TBI) has been shown to induce Mislocalization, proteolysis and aggregation of TDP-43 in humans, mice and cell culture ^53–54^, increasing the likelihood of its interaction with the pro-inflammatory GRNs.

Here, we delve deep into understanding the molecular factors that govern the interactions between GRNs and TPD-43 PrLD (termed PrLD hereafter). Our results show that extracellular GRNs-2, −3 and −5 internalize and colocalize with PrLD in the cytosol of neuroblastoma cells but do not partition into the SGs formed under stress with PrLD. In vitro biophysical experiments suggest that GRN-2 undergoes coacervation with PrLD to form liquid droplets in the oxidized state but promotes gelated/insoluble aggregates in the reduced state. With time, the droplets transform into a gelated state that shows amyloid like characteristics. Based on the empirical results derived from GRNs along with designed mutations and chemical modifications, we determine that the coacervation of GRNs and PrLD towards liquid droplet formation is predominantly driven by the negatively charged residues, while the increase in positively charged residues promotes distorted gel-like droplets and aggregation. Furthermore, we determine that the redox state of cysteines is a key modulator of phase transitions by fine-tuning LLPS or aggregation of PrLD near counterbalanced charge regimes in GRNs. Additionally, we show that RNA expunges GRNs from the PrLD droplets that provides a possible reason for not observing GRNs within the SGs.

## Results

### GRN-2 colocalizes with TDP-43 PrLD in the cytoplasm but not in stress granules

Unlike the GRNs generated in the lysosomes, the fate of those formed in the extracellular space, especially during inflammation, remains unclear although transport to the cytosol has remained a possibility ^48, 52, 55^. The ambiguity surrounding the precise localization of GRNs in pathophysiology has also hindered understanding their roles in norm and pathology ^45, 56^. We conjecture that extracellular GRNs are internalized in the neuronal cytoplasm and potentially interact with proteolytic fragments of TDP-43 modulating the latter in ALS and FTLD pathologies. To recapitulate this scenario, we utilized a simple pulse-chase assay with fluorophore-labeled GRNs ^45^. In this assay, SH-SY5Y cells transiently expressing a blue fluorescence protein tagged PrLD construct (PrLD-SBFP2) were pulsed with media containing 500 nM of HiLyte532-labeled GRN-2 after 24 h (post-transfection), and then chased for an additional hour (see Methods). The cells were then imaged under non-stress and stress (with NaAsO_2_) conditions (Fig 1). Under non-stress conditions, the transfected PrLD was observed in both the nucleus and cytoplasm (top panels; Fig 1a). Pulsing with GRN-2 showed internalization and colocalization with PrLD outside the nucleus (arrow; middle panels, Fig 1a). To see whether the colocalization was within or outside the lysosomes, as GRNs are known to be transported to lysosomes, the cells were observed using a lysosomal marker which clearly indicted GRN-PrLD puncta outside the lysosomes (arrow; bottom panels, Fig 1a). Not surprisingly, some GRN-2 was also found to be localized within the lysosomes (yellow puncta; bottom panels, Fig 1a). Fluorescence recovery after photobleaching (FRAP) of the cytoplasmic GRN-PrLD puncta showed no recovery of PrLD suggesting that they are insoluble aggregates (green, top panel; Fig 1c). In parallel, the samples were also immunodetected in the presence of TIA1 antibody, a key SG marker ^57^. In the non-stress conditions, TIA1 was located in the nucleus while PrLD was disperse throughout the cell (Fig S1a). TIA-1 was absent from the colocalized PrLD and GRN-2 puncta as would be expected under non-stress conditions (No stress; Fig S1d). Under stress conditions induced by sodium arsenite, the transfected PrLD showed puncta outside the nucleus (arrow; top panels; Fig 1b), consistent with SG formation ^11, 14^. Surprisingly, the fluidity of SGs observed by FRAP showed no recovery (orange, top panel; Fig 1c), although widely varying photo bleaching recovery rates for SGs have been reported in cellular models ^58–60^. Nevertheless, PrLD’s presence in SGs was confirmed by the colocalization of PrLD with TIA1 with a Meander’s tM1 value of 0.59 (Fig S1b). Pulsing GRN-2 onto these cells under stress showed colocalization of both the proteins as puncta in the cytoplasm and outside the lysosomes, similar to those observed under non-stress conditions (arrows; middle and bottom panels; Fig 1b). Here again, the colocalized puncta did not show fluorescence recovery (green, top panel; Fig 1c). Furthermore, GRN-2 did not show colocalization with TIA1 and PrLD suggesting that GRN-2 was not partitioned into the SGs (Fig S1d). In addition, two other GRNs known to modulate PrLD’s phase transitions ^51^, GRNs 3 and 5, also did not colocalize with TIA1 under stress conditions, but colocalized with PrLD under both non-stress and stress conditions (Fig S1d-e). Importantly, under stress, a three-way colocalization involving PrLD, GRN-2 and TIA1 was not observed. The colocalization of GRN-2 and PrLD was statistically significant with ~ 73% of the internalized GRN-2 found to be colocalized with PrLD under stress and ~ 68% under homeostatic conditions (Fig 1d). Finally, gelated material-like properties of PrLD SGs inferred by the lack of fluorescence recovery could be attributed to the absence of other domains in TDP-43. To see if this is the case, SH-SY5Y cells were transfected with the full-length wild-type TDP-43 (wtTDP43tdTomato) as a control, which showed the FRAP recovery for SGs under stress conditions (Fig S2). This possibly suggests that SG assemblies containing the truncated PrLD construct, and therefore other truncated CTFs of TDP-43 (C25 and C35), may indeed possess solid- or gel-like characteristics. However, caution needs to be exercised as a FRAP recovery alone is not an indicator of SGs as mentioned above. Taken together, the data establish that extracellular GRN-2 internalizes and colocalizes with PrLD in the cytoplasm but is not present within the SGs.

**Figure 1.**
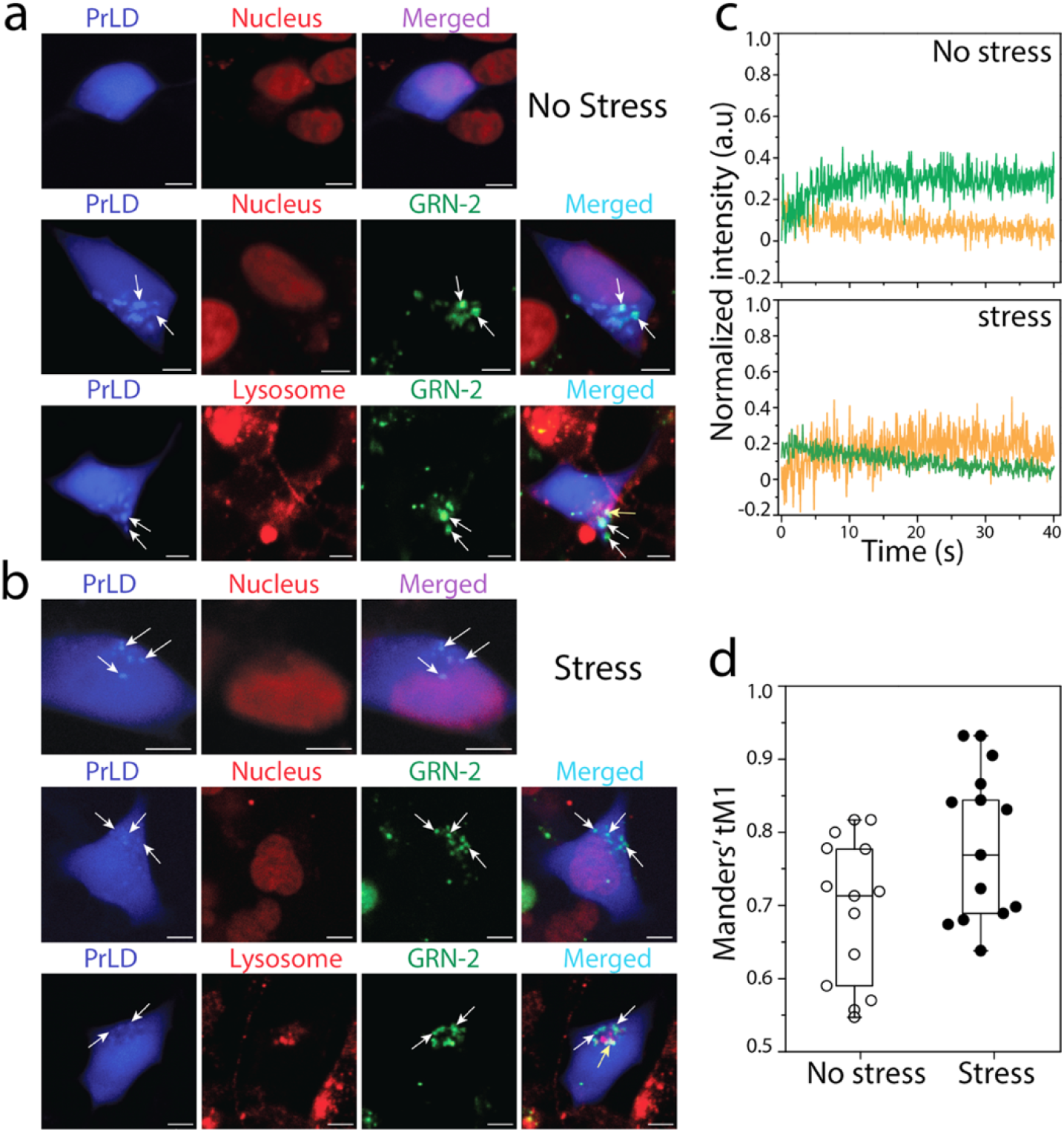
Cytoplasmic co-localization of GRN-2 and PrLD. Confocal microscopy images of live SH-SY5Ycells under homeostatic or stress conditions. a) Blue fluorescent protein tagged PrLD (PrLD-SBFP2) transiently expressed in SH-SY5Y cells alone and in the presence of Hilyte-532 labeled GRN-2 under non-stress conditions, and b) the same reactions under stress induced by sodium arsenite. White arrows indicate colocalized GRN-2 and PrLD puncta in the cytoplasm, while yellow arrows represent colocalization of GRN-2 in lysosomes (Scale bar represents 5 μm). Visualization of nucleus and lysosomes were done by staining with NucSpot^®^ Live 650 and Lysoview™ 650 respectively c) Normalized intensities of fluorescence recovery after photobleaching (FRAP) for puncta of PrLD alone 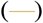 or colocalized PrLD-GRN-2 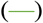 in under non-stress (top) or stress (bottom) conditions. d) Whiskers plot of Manders’ tM1 calculated for the colocalization of GRN-2 with PrLD using Fiji-ImageJ software. Each data-point represents the colocalization score of an independent cell (n = 14).

### GRN-2 modulates the phase transitions of PrLD *in vitro*

The observation of cytoplasmic puncta of colocalized PrLD and GRN-2 decoupled from SGs, prompted us to investigate their interactions in greater details. TDP-43 is known to undergo dynamic phase transitions depending on the cellular conditions to form either liquid condensates or solid aggregates ^61–63^. Understanding the molecular grammar of protein LLPS has revealed that weak, multivalent interactions between ‘stickers’ involving electrostatics, π-π (between tyrosines) or cation-π (between aromatic residues and arginines and lysines) and separated by scaffolds called ‘spacers’ are the main driving forces of condensate formation ^24, 64–65^. Disordered proteins enriched in one kind of charge will repel one another preventing self-coacervation but counter interacting charges would minimize repulsion and promote LLPS ^66^. These interactions are also observed to be the driving forces behind the self- and complex- (with RNA) coacervation of TDP-43 ^67–69^. However, the molecular determinants of heterotypic phase transitions of TDP-43 involving ligands vary depending on the partners and are still being discerned ^68^. Despite the lack of a canonical RNA binding domain, TDP-43 PrLD undergoes coacervation with RNA, which illustrates the prominence of the aforementioned interactions in phase separation ^51, 69–70^. However, the phase transitions are dictated by the balance between weak, multivalent interactions that drive LLPS, and strong, high affinity interactions that mediate gelation or solid-like phase transition often observed among amyloid proteins. Our earlier investigations into the interactions of PrLD with GRN-3 and GRN-5 showed that GRN-3 induces aggregation in both oxidizing and reducing conditions, while GRN-5 undergoes LLPS by complex-coacervation with PrLD ^51^. We conjecture that the disparity in the behavior of GRNs is due to the high net negative charge on GRN-5 (− 6; six – and no + charges, Fig 2d) as opposed to GRN-3 (− 2; seven – and two + charges, Fig 2c) that interact with PrLD (Fig 2a). It is also likely that the lack of positive charges on GRN-5 plays a role on its ability to phase separate PrLD. Here, we extended these investigations to uncouple the sequence determinants on GRNs to phase separate or co-aggregate with PrLD by investigating GRN-2 and specifically designed mutants. All three GRNs, 2, 3, and 5 possess a similar number of negatively charged acidic residues (7, 6, and 6 respectively, Fig 2b-d) but vary in the number of positively charged basic residues (2, 4, and 0, respectively, Fig 2b-d), making GRN-2 ideal to investigate. The presence of structural disorder in proteins is also known to be an implicit contributor to LLPS ^71–75^; PrLD is largely a disordered region of TDP-43 while GRN-2 is also predicted to be disordered in the fully reduced state by IUPred2A (Fig S3a-b) ^76–77^. This prediction was confirmed by biophysical characterization of GRN-2 which revealed a structure dominated by random coils in both redox-forms (Fig S4a), similar to GRNs 3, and 5 ^51^ (Fig S3c-d and S4e).

**Figure 2.**
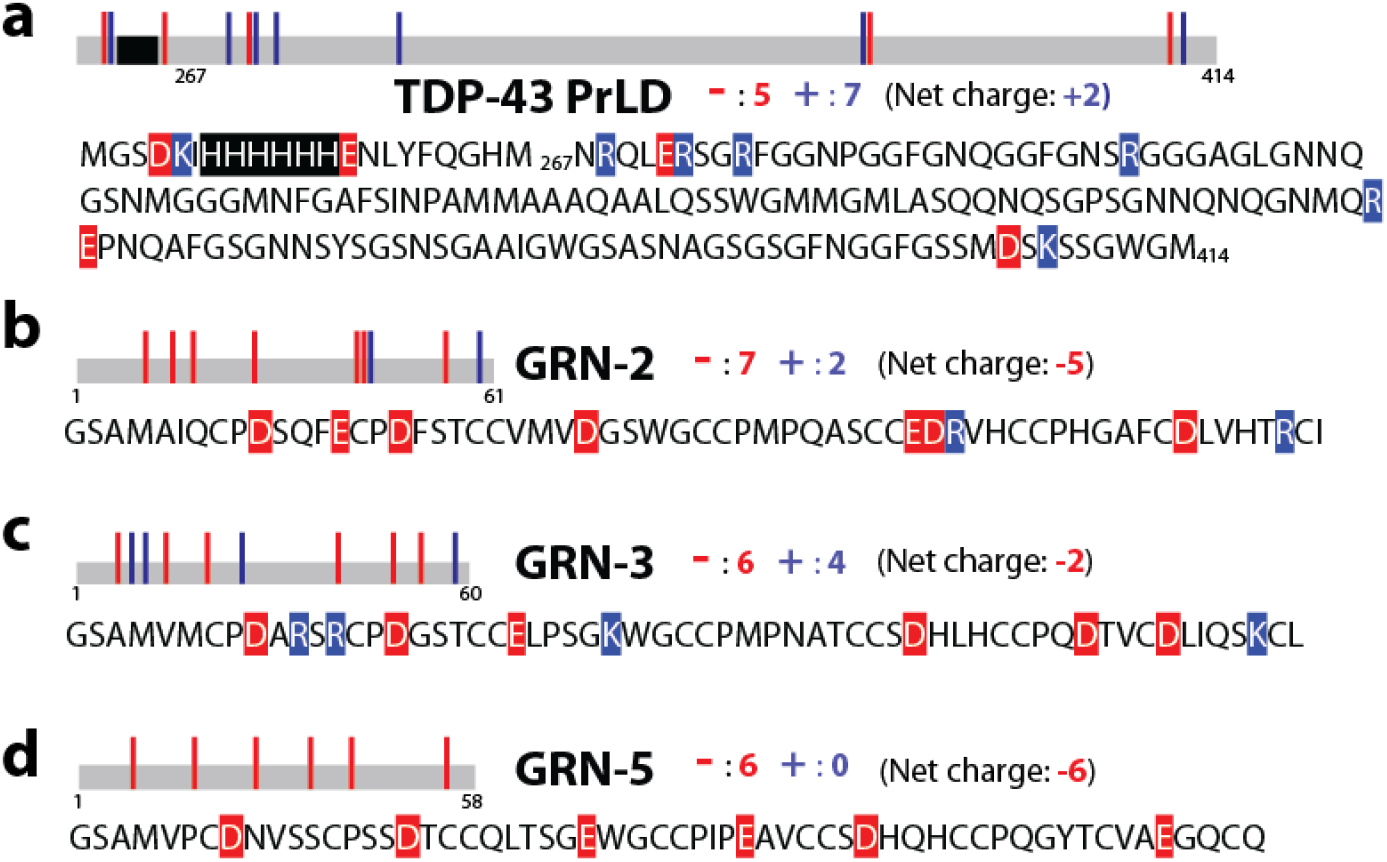
Sequences of granulins (GRNs) and TDP-43 under investigation. a) TDP-43 PrLD construct (residues 267-414) along with a hexa-histidine tag (highlighted) used in the study. b-d) Sequences of GRN-2 (b),−3 (c) and −5 (d). Sequences of proteins are annotated with acidic (red) and basic (blue) residues, along with the net charges on the respective proteins at neutral pH.

To probe the ability of GRN-2 in both redox forms to modulate phase transitions of PrLD, 40 μM GRN-2 (oxidized form) or rGRN-2 (fully reduced form) were mixed with 20 μM PrLD buffered in 20 mM MES, pH 6.0 and the samples were visualized by differential interference contrast (DIC) microscopy (Fig 3a-c). Immediately upon coincubation of GRN-2 and PrLD, the sample phase separated into liquid droplets ranging between 2 and 12 μm diameter (Fig 3a). The droplets also showed classic wetting behavior suggestive of their liquid-like characteristics (arrow; Fig 3b). The colocalization of proteins within the droplets was confirmed using orthogonal fluorophores, HiLyte405 (blue) and HiLyte647 (red) for GRN-2 and PrLD, respectively, and their liquid-like properties by coalescence and fusion of droplets (Fig 3b). In stark contrast, samples containing rGRN-2 and PrLD showed the formation of solid insoluble aggregates (Fig 3c) which also involved colocalization of both proteins (Fig 3d). These results indicate that GRN-2 in the oxidized form induced LLPS of PrLD, while the reduced form induced gelation/aggregation. primarily by Then LLPS/aggregation phase boundaries were determined for GRN-2 and rGRN-2 with PrLD using turbidimetric assays (Fig 3e-g). In all these experiments, LLPS and aggregation were differentiated based on microscopic examination of the samples and phase boundaries established by considering an OD_600_ value of 0.140 a.u. as a cut-off for phase transitions (see Methods). To see if the negative charges drive GRN-PrLD LLPS, phase separation was observed at two different pH; at pH 6.0 where the acidic residues have a net negative charge and at pH 4.5 where the acidic residues are close to being neutral. At pH 4.5, no phase separation was observed for GRN-2 or rGRN-2 with PrLD (Fig 3e, pH 4.5) but at pH 6.0, the proteins underwent phase transitions uniformly at a concentration of 10 μM GRN-2 or rGRN-2 in all PrLD concentrations tested (Fig 3e, pH 6.0). However, in stark contrast, under fully reducing conditions, co-incubations of rGRN-2 with PrLD showed gelation/aggregation. The phase diagrams of GRN-3, GRN-5 with PrLD reveal both similarity and disparity in their phase transitions as previously observed ^51^(Fig 3f-g); In an acidic environment, neither GRN-3 nor GRN-5 in both redox conditions undergoes phase transition with PrLD (Fig 3f) similar to GRN-2. At pH 6.0, both oxidized and reduced forms of GRN-3 show gelation/aggregation exclusively, albeit with a delay (solid precipitates observed after ~24 h), while incubation with GRN-5 showed LLPS exclusively (Fig 3g) ^51^. Together, these results unambiguously indicate that the negatively charged residues on GRNs (evident from GRN-5, and data from assays at pH 4.5) are significant contributors of the coacervation with PrLD while an increasing number of positive charges on the protein modulates the dynamics towards gelation/aggregation (as observed from GRN-2 and GRN-3). Next, we questioned whether GRN-PrLD coacervation was driven by electrostatic interactions, in which case LLPS will be sensitive to salt concentrations. To investigate this, turbidities of different stoichiometric incubations of PrLD and GRN-2 with increasing NaCl concentrations were measured. The data showed progressive decrease in turbidity values with increasing salt concentrations likely due to disruption of electrostatic interactions by NaCl and thus confirming electrostatically driven phase separation between GRN-2 and PrLD (Fig 3h). We also investigated the potential role of hydrophobic interactions in mediating LLPS by treating the condensates of GRN-2-PrLD with 1,6-hexanediol (HD), an agent that is known to disrupt weak hydrophobic interactions and dissolve droplets ^78–79^. Post treatment analysis showed persistence of the condensates even after treatment with 10% HD (top panels; Fig 3j) as evidenced by turbidity, albeit with an ~ 45% reduction in their levels compared to pre-HD treated sample. This suggests that hydrophobic interactions contribute to LLPS to some degree. Similar treatments of HD to PrLD-RNA coacervates showed a ~15 % decrease in turbidity reaffirming that LLPS involving the two was driven mainly by electrostatic interactions (middle panels; Fig 3j). In contrast, homotypic coacervation of PrLD induced by salt (300 mM) showed dissolution as much as 90% of droplets upon HD treatment (bottom panels; Fig 3j) suggesting that the salt-induced phase separation is predominantly driven by hydrophobic interactions as proposed earlier^80^. Together, the data suggests that GRN-2-PrLD coacervation is an electrostatically driven process. Finally, the temperature dependence of PrLD-GRN-2 phase transitions was examined to establish the upper critical solution temperature (UCST) ^69, 81^. As expected PrLD underwent self-coacervation in no salt conditions at 4 °C that was attenuated at 25 °C suggesting a UCST between these temperatures (Fig 3k). Co-incubation with GRN-2 increased UCST to > 49 °C, while the sample with rGRN-2 showed gelation/aggregation in all the temperature range recorded (Fig 3k).

**Figure 3.**
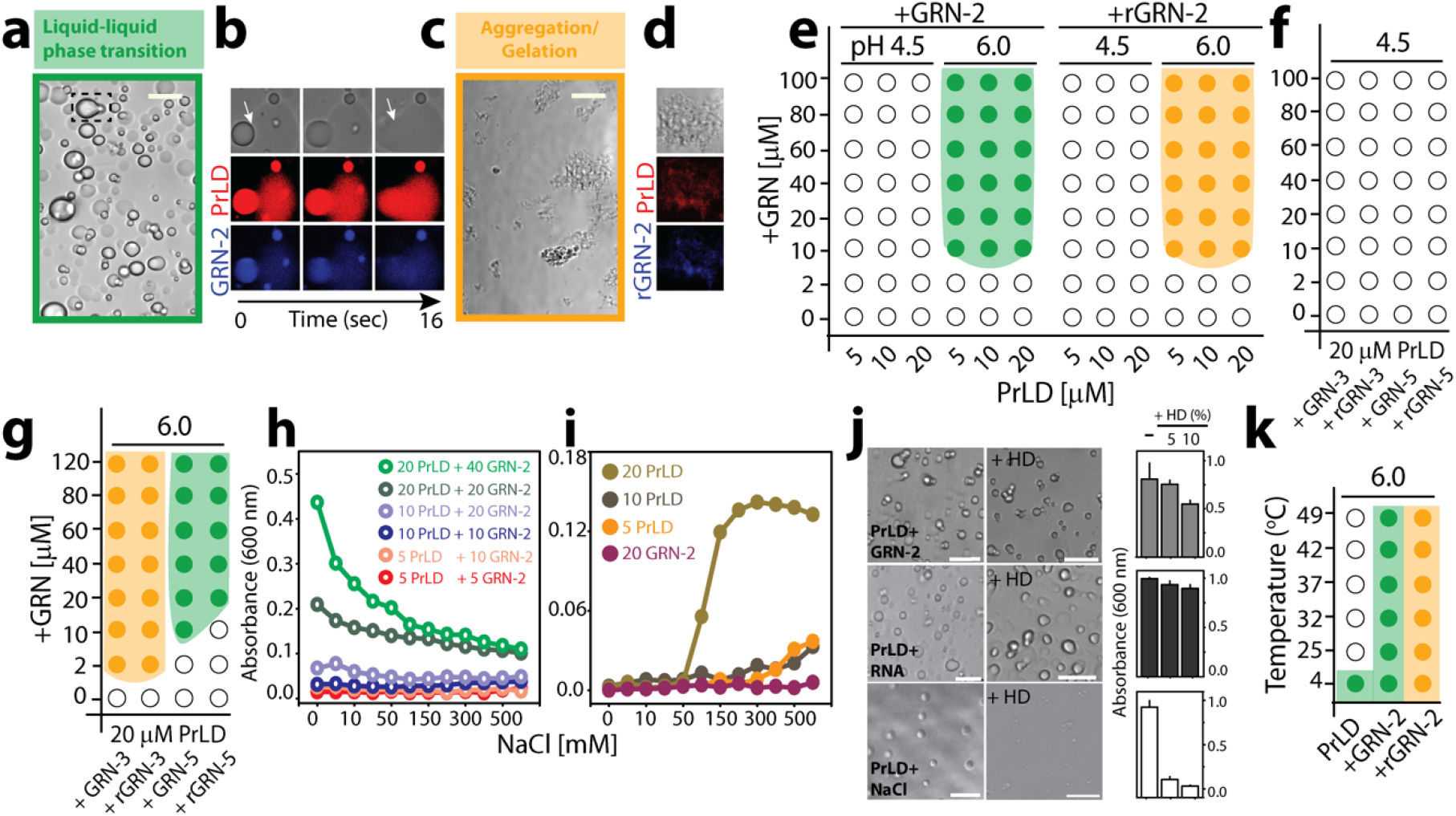
Phase transitions involved in GRN–TDP-43 PrLD interactions. a-d). DIC microscopy images of a mixture containing 20 μM PrLD with 40 μM fully oxidized GRN-2 (a-b) or reduced rGRN-2 (c-d) in 20 mM MES, pH 6.0. GRN-2 undergoes LLPS with PrLD (highlighted with green box) that shows the coalescence of droplets (a, dashed box) while rGRN-2 undergoes gelation/aggregation (saffron box). GRN-2 or rGRN-2 labelled with HiLyte 405 (blue) and PrLD with HiLyte 647(red) were visualized using confocal fluorescence microscopy which shows fusion of droplets for GRN-2 (white arrow; b) and deposition of solid aggregates for rGRN-2 (d). e)Phase diagram for GRN-2 or rGRN-2 with PrLD generated by varying the pH (buffered with 20 mM ammonium formate at pH 4.5 or 20 mM MES at pH 6.0) and concentrations of the respective proteins. g-h) Phase diagram for varying concentrations of GRN-3 and GRN-5 with 20 μM PrLD at pH 4.5 (f) and pH 6.0 (g). (h) Turbidity measurements monitored at 600 nm for PrLD-GRN-2 reactions with different stoichiometries in increasing NaCl concentrations (1-750 mM). (i) Turbidites of the control reactions for samples in (h). j) A mixture containing 40 μM GRN-2 with 20 μM PrLD was treated with 5 and 10 % 1,6-hexanediol (HD) to monitor its effect on the LLPS of the proteins which was quantified using OD_600_ values pre- and post-HD treatment. Similar treatment was performed for condensates of 20 μM PrLD with either RNA (75 μg/mL) or NaCl (300 mM). The turbidity values were normalized with respect to their values pre-HD treatment (n = 3). The concentrations of components used in this assay (GRN-2, RNA, PrLD and NaCl) were carefully chosen to be well within the demixing boundary conditions of the individual phase transitions. k) The turbidity of 20 μM PrLD alone, or in presence of 40 μM GRN-2 or rGRN-2 measured as a function of temperature. The phase boundaries were established by considering an OD_600_ value of 0.14 as a cut-off for phase transitions. Scale bar represents 10 μm. For e-k, three independent data points (n=3) were averaged.

### Negatively charged residues in GRNs drive LLPS while the positively charged ones enhance gelation or aggregation

The pH dependence on phase transitions (Fig 3e-f) and HD treatment (Fig 3j) provided important clues regarding the involvement of electrostatic interactions, especially the negatively charged residues. To further investigate these in detail, the phase behavior of the three granulins, GRN-2, −3, and −5 along with specific mutants of GRN-2 were investigated. As mentioned above, all three GRNs are isoelectronic with respect to the negative charges (−7, −6, and −6, respectively) but contain a varying number of positively charged residues (2, 4 and 0, respectively) (Fig 2). As previously shown ^51^, microscopic investigations of GRN-3 incubated with PrLD in both redox states showed aggregation (saffron box) while GRN-5 showed LLPS (green box) under both redox conditions (Fig 4a and b). FRAP confirmed the observations with a rapid recovery for GRN-5 (liquid) and attenuated recovery for GRN-3 (solid) (Fig 4a and b). On the other hand, GRN-2 showed LLPS in the oxidized state and aggregation in the reduced state (Fig 4c). This suggests that the number of positive charges in GRN-2 is enough to mitigate LLPS in the reduced state but not enough to attenuate LLPS and promote aggregation in the oxidized state. We conjectured that while negative charges on the GRNs promote LLPS, the positively charges residues tend to shift the equilibrium towards aggregation and solid aggregates. To further substantiate the role of positive charges, a construct was generated in which the two positive charges on GRN-2 were mutated to glutamate (negatively charged) residues to generate GRN-2 (+/-) mutant that was devoid of any positive charges (net −9 charge), which rendered the mutant similar to GRN-5 electronically. If our hypothesis was correct, abrogation of all positively charged residues would promote LLPS in both oxidized and reduced states identical to GRN-5. Co-incubation of GRN-2 (+/-) and PrLD did show formation of liquid droplets in both fully oxidized and reduced conditions (Fig 4d), further cementing our hypothesis that negative charges drive LLPS. To further illustrate this, GRN-7, which has the same number of acidic (−6) residues as the other GRNs but has high number of basic residues (+8), with an overall positive charge (+2), was co-incubated with PrLD (Fig S5a). Based on the on the inferences drawn thus far, such enrichment of positively charged residues will drive aggregation of PrLD, which was confirmed by the presence of solid aggregates on the co-incubated sample monitored for 36 h (Fig S5c). Unfortunately, a deeper investigation on GRN-7 was precluded by the difficulties in obtaining a pure protein that was prone to substantial proteolytic degradation over time (data not shown). Lastly, we evaluated the droplets formed by these proteins with PrLD (Fig S8). The droplet area (μm^2^) of GRN-2 with PrLD shows a normal distribution centered at ~25 μm^2^ (Fig S8, gray), which is similar to that observed for GRN-2(+/-) with PrLD (Fig S8, red). On the other hand, droplets formed in the reaction of rGRN-2(+/-) with PrLD shows noticeable reduction in droplet areas with a normal distribution centered at ~10 μm^2^, suggesting a diminished ability of the reduced form to undergo LLPS. Furthermore, to ascertain the contribution of hydrophobic aromatic residues in LLPS, F, Y and W residues of GRN-2 were mutated to A to generate GRN-2ΔAro mutant. Unfortunately, significant degree of thiol oxidation to sulfinic acid was observed when expressed (Fig S9a) possibly due to more exposed thiols with lack of hydrophobic core, which rendered the results indiscernible. Co-incubation of GRN-2ΔAro with PrLD showed amorphous deposits with no recovery observed in FRAP suggesting solid nature of the deposits (Fig S9b). As sulfinic acid imposes a strong hydrophilic effect (elaborated further below), precise effects of the lack of aromatic residues were undeterminable.

**Figure 4.**
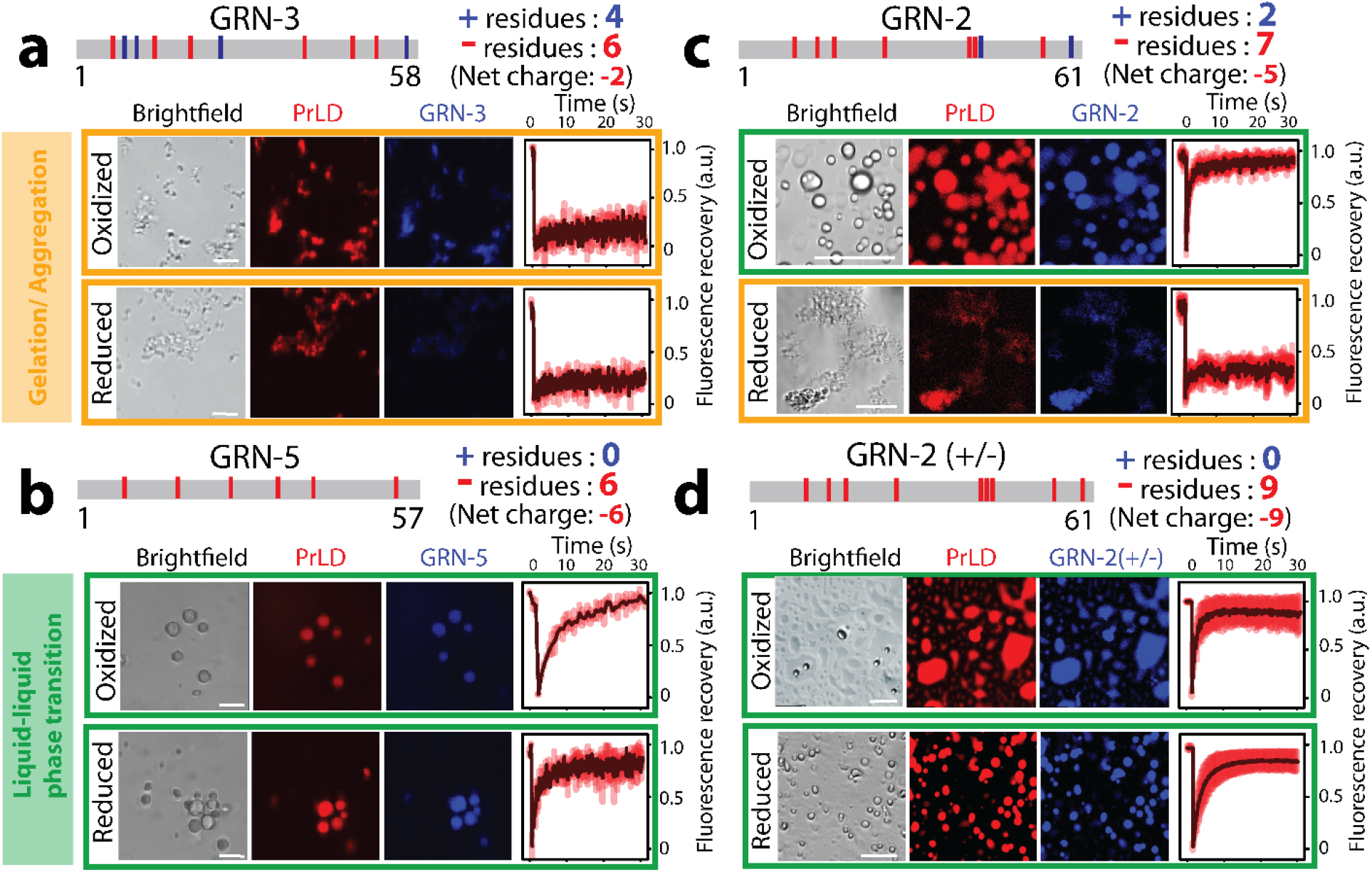
Phase transitions of GRNs and specific mutants. a-d) Sequence of GRN-3, GRN-5, GRN-2 and GRN-2(+/-) annotated with negative and positive charges present and the net charges on the protein at neutral pH (top of each panel). Individual samples are generated by mixing 40 μM of the respective GRN with 20 μM PrLD, along with 1% (molar) fluorophore-labeled proteins (GRNs are labeled with HiLyte-405 and PrLD with HiLyte 647) buffered in 20 mM MES, pH 6.0. Samples were visualized by fluorescence microscope and their internal dynamics were analyzed by FRAP. Individual micrographs are highlighted with a green border for LLPT and a saffron border for LSPT. The fluorescence recovery curves were normalized based on pre-bleaching fluorescence intensities. Scale bar represents 10 μm. The images are representatives from at least three independent data points (n=3).

### The redox state of cysteines in GRNs further tunes phase behavior of PrLD coacervates

It is now clear that the coacervation of GRN and PrLD is driven predominantly by electrostatic interactions; while negative charges drive LLPS the increase in the number of positive charges shift the equilibrium towards aggregation. However, what role does the oxidation state of thiols play in this process remained unclear. Therefore, to understand their effect, microscopic analysis was performed on the co-incubations of GRN-2 and PrLD as a function of a reducing agent (Fig 4). To mimic local redox fluxes within the cellular cytoplasm ^82^, the reactions containing 40 μM GRN-2 and 20 μM PrLD with increasing concentrations (50-600 μM) of the reducing agent tris(2-carboxyethyl)phosphine (TCEP) were probed by fluorescence microscopy (Fig 4a). The TCEP concentrations used generate partially reduced (50-300 μM) to completely reduced (600 μM) thiols in GRN-2. In the absence of the reducing agent, GRN-2, as expected, underwent LLPS with PrLD forming liquid droplets that undergo fusion and coalescence as observed previously (0 μM TCEP; Fig 5a). To ascertain the droplets’ fluid-like properties, their internal dynamics were analyzed by FRAP which showed a rapid recovery for PrLD reiterating an archetypal fluidic characteristic of the droplets (red curve, 0 μM TCEP; Fig 5a). For reasons not clear at this time, attenuated mobility for the colocalized GRN-2 within the liquid droplets was observed as seen in its recovery curve, (blue curve, 0 μM TCEP; Fig 5a). The observed LLPS by oxidized GRN-2 could also arise from the network of intra-molecular disulfides bonds that limit the solvent accessibility of some of the stickers in the sequence. This, in part is supported by the attenuated FRAP recovery rates for GRN as opposed to PrLD (Fig 5a and b), and the solvent exposure of the single tryptophan residue in the middle of the sequence in oxidized and reduced states (Fig S10).

**Figure 5.**
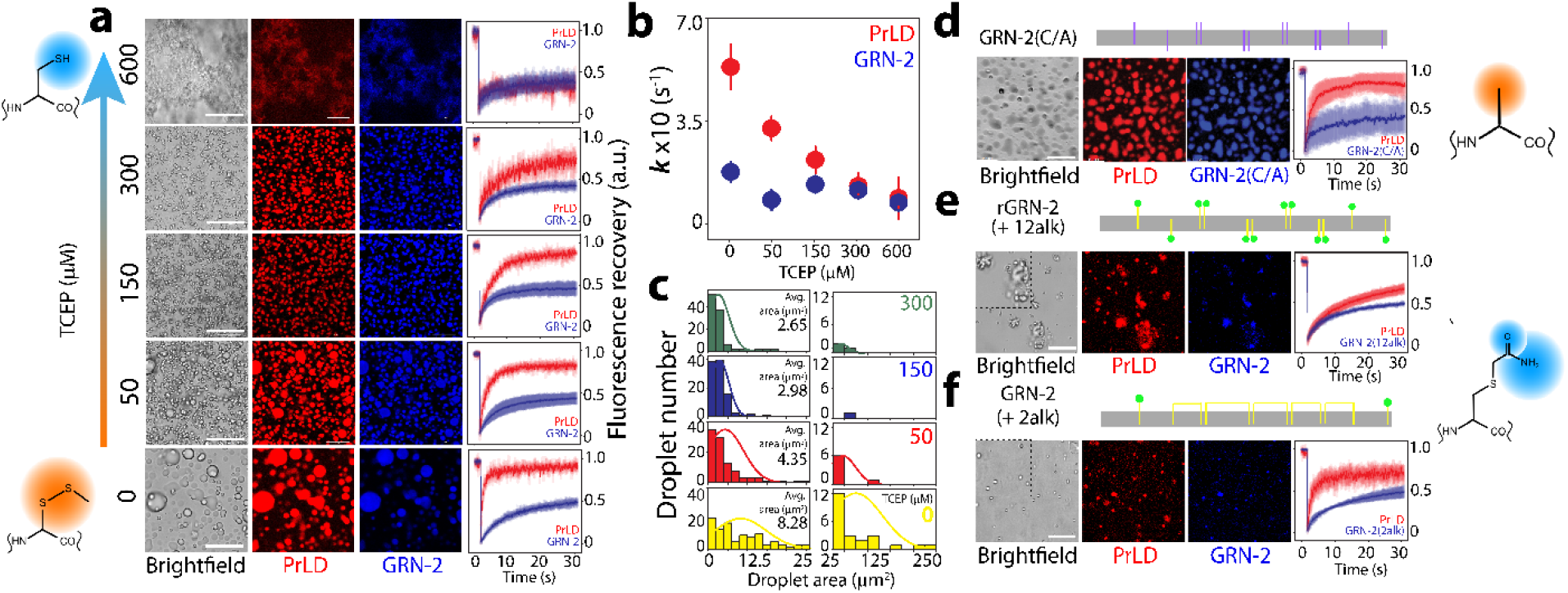
Redox state of GRN-2 fine tunes the phase transitions with PrLD. a) Reactions containing 20 μM PrLD with 40 μM GRN-2 were initiated separately with increasing TCEP concentrations (50-600 μM) buffered in 20 mM MES, pH 6.0. The hydrophobic and hydrophilic environments around oxidized and reduced cystines, respectively, are schematically presented. b) Samples were visualized by fluorescence microscopy and probed with FRAP recovery (n = 3). Individual recovery rates of PrLD (red) and GRN-2 (blue) in the reactions of a) obtained using the initital rate method (n = 3; detailed in Materials and Methods). c) Distributions of droplet area (μm^2^) observed within the micrographs of reactions in (a). The droplet area distributions were extracted using ImageJ where a total of 100 droplets was considered for each sample and Normal distribution are plotted (detailed in Materials and Methods). d) Schematic depicting the GRN-2(C/A) mutant (cysteines replaced with alanines; purple vertical bars). The micrographs represent reactions of 20 μM PrLD with 40 μM GRN-2(C/A) buffered in 20 mM MES, pH 6.0 along with the FRAP analysis. e-f) Micrographs of mixtures containing 40 μM alkylated-GRN-2 (e; 2-free thiols alkylated, f; 12-free thiols alkylated) generated using iodoacetamide, with 20 μM PrLD along with the FRAP analysis. For visualization using fluorescence microscopy, all samples contained 1% fluorophore-labeled proteins (GRN-2, GRN-2(C/A), and alkylated forms were labeled with HiLyte-405 while PrLD was labeled with HiLyte-647). All reactions were initiated and imaged at room temperature. Scale bar represents 20 μm.

In the presence of 50 μM TCEP (partially reducing conditions), the magnitude of droplet formation was dampened and structures with an altered morphology were visible alongside spherical droplets (50 μM TCEP; Fig 5a). Increasing concentrations of TCEP (150-600 μM) showed numerous smaller droplets and a progressive decrease in fluorescence recovery rates for PrLD (Fig 5a and 5b). The rate constant (*k*) deduced by an initial rate method from the FRAP data showed a first-order kinetics for PrLD recovery as a function of reducing agent concentrations, while GRN-2 showed a diminished rate of recovery in fully oxidized and partially reduced (50 μM TCEP) conditions (Fig 5b). In fully reducing conditions, the mixture of rGRN-2 and PrLD underwent aggregation, as observed previously (Figs 3 and 4) generating solid, fibril-like aggregates and was devoid of any droplets, as confirmed via microscopy and FRAP data (600 μM TCEP; Fig 5a). The use of other reducing agents such as dithiothreitol (DTT) and glutathione (GSH) also showed similar results (Fig S6). Furthermore, analyses of droplet size as a function of the reducing agent provided some interesting insights (Fig 5c). The droplet size distributions, in terms of the surface area, were clubbed into two categories of small (area < 25 μm^2^) and large droplets (25 < area < 250 μm^2^). The coacervates in fully oxidized conditions showed a wide distribution of droplet sizes. The surface areas of small droplets were centered at ~10 μm^2^, with an average droplet size of 8.2 μm^2^ (yellow; Fig 5c). In addition, a notable number of droplets larger than 25 μm^2^ were also observed in the sample with an average surface area of ~ 60-70 μm^2^ (yellow; Fig 5c). Increasing TCEP concentrations (50-600 μM) revealed a discernible shift towards smaller droplet distributions with average surface areas of 4.35, 2.89 and 2.65 μm^2^ for 50, 150, and 300 μM TCEP concentrations, respectively (Fig 5c). We attribute this behavior to the reduction in surface tension introduced by way of thiols solvation or hydrogen bonding interactions.

These results bring out two important points; first, the modulation of phase transitions by potential redox fluctuations does not seem to be abrupt but requires a complete reduction of cysteines to promote gelation or aggregation. Second, cysteine thiols seem to play a role in shifting the equilibrium from the liquid to the gelated/aggregated phase possibly due to a hydrophilic environment generated and by engaging in hydrogen bonding interactions with bulk water (solvation) and/or with PrLD (Fig 5a). If this is the case, we argued that abrogation of cysteines (and therefore, thiols) will render GRN-2 to behave as in fully oxidizing conditions. To test this, all 12 cysteines in GRN-2 were mutated to alanines to generate a GRN-2(C/A) mutant. Indeed, co-incubation of PrLD with GRN-2(C/A) showed the formation of liquid droplets with rapid fluorescence recovery confirming a fluid-like character (Fig 5d). The similarity in the behavior of GRN-2(C/A) and GRN-2 is also apparent from the droplet size observed (Fig S8). The droplet areas of the mutant form (Fig S8, green) have a similar distribution to that of the wild type (Fig S8, gray), with an average value of ~25 μm^2^. On the contrary, when all the free thiols in the fully reduced form of rGRN-2 were alkylated and capped with acetamide (GRN-12alk), a moiety, which is expected to be both solvated by bulk water and engage in hydrogen bonding interactions with PrLD perhaps to a greater extent than thiols, the coacervates formed solid like structures with attenuated internal mobility as revealed by FRAP (Fig 5e). Similarly, acetamide alkylation of two free thiols present in the fully oxidized samples (Fig S4), which mimics partially reducing conditions also showed no change from those incubated in the presence of 50 μM TECP (Fig 5f). The effect of solvent interactions with thiols in the reduced state was further supported by our serendipitous discovery with GRN-2ΔAro mutant. Despite stringent experimental precautions to prevent oxidation of thiols during expression and purification, GRN-2ΔAro mutant showed up to five cystine thiols were oxidized to sulfinic acids (Fig S9a). Since sulfinic acids are strongly hydrophilic, they are likely to be solvated and engage in hydrogen bonding interactions with water. Thus, one could expect aggregation with GRN-2ΔAro mutant when co-incubated with PrLD, which is exactly what was observed (Fig S9b). Together, the data bring to light the role of cysteine in phase transitions of PrLD and GRNs, which is one of modulatory nature, fine-tuning the transitions only in GRN-2, which happens to fall in category of partially counter-balanced positive and negative charges. As observed before ^51^, GRNs 3 and 5 showed no effect on respective phase transitions due to the extremity of the electrostatics in their respective cases.

### GRN-2 delays but induces ThT-positive aggregates of TDP-43 PrLD in oxidized or reduced states

We have previously shown that the concentration-dependent LLPS of PrLD by GRN-5 or aggregation by GRN-3 are accompanied by a delayed emergence of thioflavin-T (ThT) fluorescence ^51^, which is a reporter of amyloid-like aggregates ^83^. To see whether GRN-2 in both redox forms induces ThT-positive PrLD aggregates, the co-incubations of increasing concentrations of GRN-2 or rGRN-2 and PrLD were monitored for 24 hours in the presence of ThT (Fig 6a). All reactions showed slightly elevated ThT levels at the beginning (0h; Fig 6a) but failed to show sigmoidal increase of ThT fluorescence as observed for the control PrLD (Fig 6a). This observation was similar to that observed previously with GRN-3 and GRN-5 ^51^. Microscopic investigations of the reactions with GRN-2 showed the formation of liquid droplets in all stoichiometric equivalence immediately after incubation (0h; Fig 6b). After 24h, the droplets continue to be present although a few of them showed morphological changes suggesting possible gelation (24h; Fig 6b). To probe this possibility, we performed FRAP on the structures after a 24h period. Compared to the fluorescence recovery shown by the liquid droplets of 40 μM GRN-2 with 20 μM PrLD (Fig 6b, FRAP, 0 h; red), the distorted droplets in all stoichiometric reactions show attenuation in their recovery kinetics (Fig 6b, FRAP, 24 h; brown) confirming the progressive ‘gelation’ of the liquid droplets. In addition, this attenuation positively correlates with GRN-2 concentrations (Fig 6b). On the other hand, under fully reducing conditions, incubations of rGRN-2 with PrLD showed an instantaneous formation of insoluble aggregates that persisted after 24 h (Fig 6c). However, after > 60 hours of incubations with oxidized and reduced GRN-2 in lower stoichiometric equivalence showed increases in ThT intensities towards amyloid like aggregates (Fig S6).

**Figure 6.**
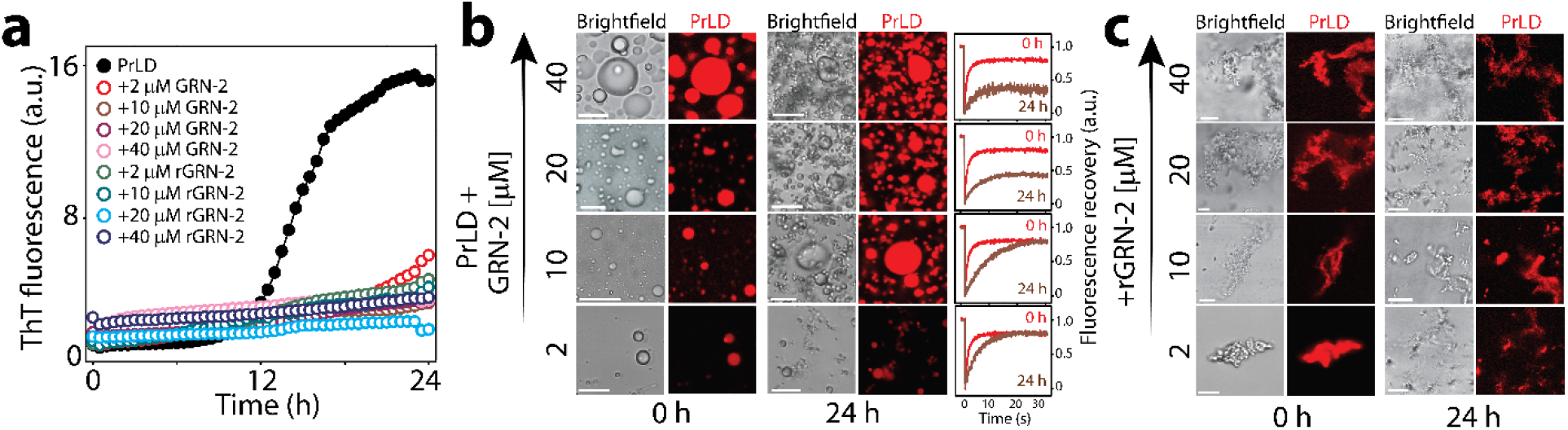
Amyloid formation of PrLD in presence of GRN-2. a) The amyloid formation of 20 μM TDP-43 PrLD alone (●) or in presence of varying concentrations (2-40 μM) of GRN-2 or rGRN-2, buffered in 20 mM MES, pH 6.0, monitored using 15 μM ThT for a period of 24 h at 37 °C under quiescent conditions. b-c) Micrographs of reactions in (a) visualized using brightfield and fluorescence microscope with 1% fluorophore-labeled PrLD (HiLyte-647) imaged at 0 h and after an incubation of 24 h. b) The droplets observed initially upon mixing GRN-2 with 20 μM PrLD were subjected to FRAP analysis immediately and after 24 h (0 and 24 h, red and brown curves, respectively). c) Microscopic visualization of PrLD in the presence of varying concentrations of rGRN-2 (2-40 μM) shows the formation of solid deposits at 0 h and 24 h. Scale bar represents 10 μm.

### RNA competes and displaces GRN-2 from PrLD coacervates

Under cellular stress conditions, PrLD transfected in SH-SY5Y cells is either found in SGs along with TIA1 or colocalized with GRNs in the cytoplasm, but colocalization of all three was not observed (Fig S1). We conjectured that it is due to potential competition between GRNs and RNA, in which the latter displaces the former in SGs. To test this, ternary interactions between RNA, GRN-2, and PrLD were investigated in vitro (Fig 7). PrLD (20 μM) was co-incubated with three GRN-2 concentrations of 10, 20 and 40 μM, and each co-incubated sample was investigated with the addition of increasing RNA (poly A) concentrations (0-150 μg/mL). The positive control, PrLD, and RNA showed prototypical turbidity increase with increasing RNA concentrations that saturated about 50 μg/mL and showed a slight drop in turbidity at 150 μg/mL (●; Fig 7a). Increasing RNA concentrations in PrLD-GRN-2 co-incubations also showed similar trends but with higher initial turbidity values (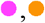, and 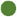; Fig 7a). Irrespective of the concentration of GRN-2 in the mixture, the eventual turbidity values of all samples converge to that shown by PrLD with 150 μg/mL RNA. This hints at the similar compositional make-up of the final condensates which is insensitive to GRN-2 concentration in the presence of high RNA concentration (Fig 7a). To probe the internal dynamics of these interactions, we used fluorescence microscopy with fluorophore-labeled proteins and repeated the experimental setup. Co-incubations of varying concentrations of GRN-2 with PrLD, in the absence of RNA, showed droplet formation as expected (0 RNA; Fig 7b-e). The addition of increasing amount of RNA to 10 μM GRN-2 incubations showed a steady displacement of GRN-2 (blue) from within the droplet from ~ 50-75 μg/mL, while PrLD (red) remained largely unaffected (Fig 7c). Between 100 and 150 μg/mL of RNA, GRN-2 has been completely displaced from the droplet (Fig 7c). Interestingly, the displacement and substitution of GRN-2 did not occur concertedly but involved the formation of coexisting multiphasic condensates with coacervates of PrLD-RNA engulfing those of PrLD-GRN-2 as deduced by the fluorescence intensity plots (Fig 7c).

**Figure 7.**
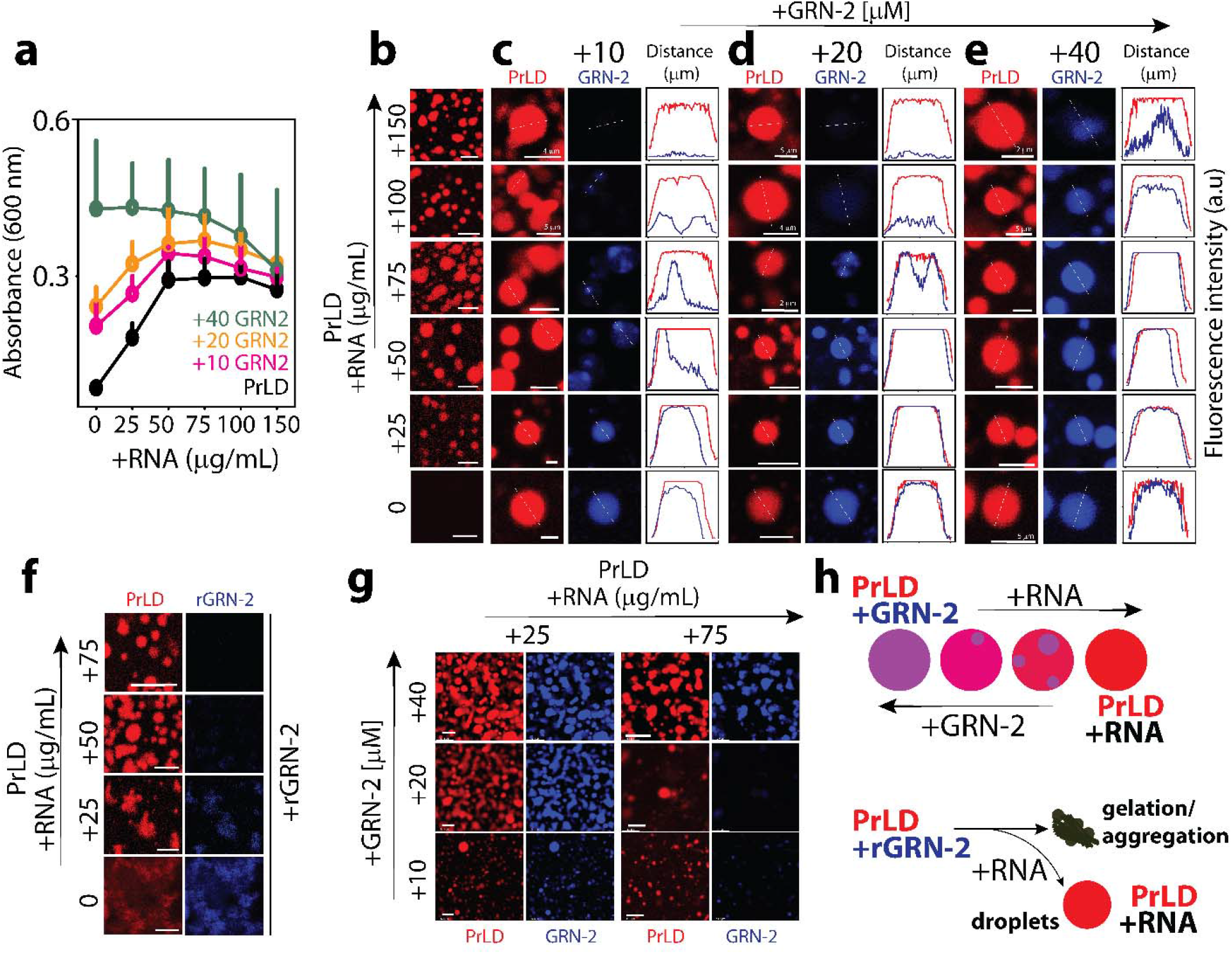
Ternary interactions between GRN-2, PrLD and RNA. a) Turbidity plots obtained by titrating 20 μM PrLD with increasing RNA concentration (25-150 μg/mL) in the presence (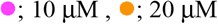 or 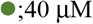 GRN-2) or the absence of GRN-2 (●) (n =3). b-e) Confocal micrographs depicting selective droplets observed in reactions from (a). Displacement of GRN-2 upon titration with RNA was observed by the fluorescence intensity profiles of the respective fluorophore-labeled proteins (GRN-2 with HiLyte-405 and PrLD with HiLyte-647) was extracted across the width of the droplets using ImageJ (detailed in Materials and Methods). f) Microscopic visualization of complex coacervates of 20 μM PrLD with varying RNA concentrations (25-75 μg/mL) generated initially were titrated with 40 μM rGRN-2. g) Condensates of 20 μM PrLD with varying RNA concentration (25 or 75 μg/mL) were titrated with increasing GRN-2 concentration (10-40 μM) and were visualized microscopically. h) Schematic depicting the outcome of ternary interactions amongst GRN-2, RNA and PrLD. All reactions were intiated and imaged at room temperature. Unless specified, scale bar represents 10 μm.

A similar behavior was observed with samples containing increased GRN-2 concentrations (20 and 40 μM) also (Fig 7d-e). Such co-existing heterotypic condensates have also been observed in ternary complexes involved in prion-like polypeptide, RNA and arginine-rich polypeptides ^84^. Similar to Kaur and colleagues, we observe a ternary system in which RNA-PrLD interactions out compete and dominate PrLD-GRN-2 interactions by displacing GRN-2. This also suggests that GRN-2 and RNA are mutually exclusive binding partners for PrLD demixing and LLPS, which in turn explains why GRN-2 are not observed in SGs. Furthermore, we also examined the effect of rGRN-2 on the condensates of PrLD-RNA (Fig 7f). Here, condensates of 20 μM PrLD with varying RNA concentrations (25-75 μg/mL) were generated and 40 μM rGRN-2 was added followed by visualization using fluorescence microscopy. We observed that at a low RNA concentration (25 μg/mL), rGRN-2 is able to interact with unengaged PrLD within the mixture and drive its aggregation (Fig 7f, +25 RNA) similar to that observed in the absence of any RNA (Fig 7f, 0 RNA). In the condensates of PrLD-RNA generated with higher RNA concentrations (50-75 μg/mL), the external addition of rGRN-2 leads to minimal or no aggregation with the sustenance of RNA-PrLD condensates, as seen in the micrographs (Fig 7f, +50 and +75 RNA). We also performed an inverse experiment where GRN-2 was titrated on PrLD-RNA droplets (Fig 7g), and observed that high concentrations of GRN-2 coacervates with PrLD presumably displacing RNA. These results provide potential clue to the absence of GRNs in SGs, which is likely the result of high amounts of RNA (and possibly other proteins) in SGs displacing GRNs (Fig 7h).

## DISCUSSION

The precise role of GRNs in the pathophysiology of neurodegenerative disorders has remained an open question. In general, under what conditions and cellular cues are they generated from their percussor PGRN, and what role do they play in cellular functions and dysfunction has been a point of debate ^44–45, 85–88^. What we do know is that PGRN secreted from microglia and astrocytes is transported into neuronal lysosomes by a sortilin-mediated pathway ^88–89^ and that its proteolytic processing generates GRNs within the lysosomes ^45, 90^ but what functional roles they play in lysosomes remain uncertain. PGRN also undergoes proteolytic cleavage extracellularly during chronic inflammation ^42^ such as those during neurogenerative disease, and the fate of extracellular GRNs on neuronal functions also remains completely unknown. Here we investigated the cellular localization and interaction of extracellular GRNs (GRN-2, −3 and −5) with TDP-43 by treating them onto SHSY5Y neuroblastoma cells that transiently express TDP-43 PrLD. These studies along with in vitro biophysical investigations on the interactions between the two proteins in greater detail have brought out some unexpected properties of GRNs and PrLD that unravel GRNs’ potential role in FTLD and related pathologies.

To understand the fate of extracellular GRNs on neurons, we labeled GRNs only with organic fluorophores (HiLyte) to visualize their uptake but were devoid of any tags that may predispose them for specific cellular localization. The data indicate that the GRNs are taken up by SH-SY5Y cells to be localized not only within lysosomes, which is well-known, but in the cytosol also. More importantly, GRNs colocalize with TDP-43 PrLD in the cytoplasm both under stress and non-stress conditions indicating their potential involvement in FTLD and ALS pathologies (Fig 1). Furthermore, under stress conditions, based on the lack of FRAP recovery and absence of colocalization with the SG marker TIA1 (Fig S1), we could conclude that GRNs fail to partition within the SGs. In addition, we infer that the cytosolic colocalizations observed are complex cytoplasmic coacervates of PrLD and GRNs. Interestingly, unlike with PrLD, GRNs failed to colocalize with full-length TDP-43 either in non-stress conditions or within SGs under stress conditions (Fig S2), illustrating that GRNs may specifically interact with the pathogenic proteolytic fragments of TDP-43, for which the PrLD constitutes the major part ^91^, further implicating their role in pathology.

The dynamics of interactions between GRN-2 and PrLD investigated here have furthered our understanding of the possible mechanisms by which GRNs could modulate intracellular inclusions of TDP-43. Coacervation of TDP-43 with RNA has been known to be electrostatically driven ^68, 70^. In the present study, we have used PrLD containing an N-terminal hexa-histidine tag (Fig 2a). We observed that cleavage of the tag prohibitively decreases the yields of the protein for biophysical investigations as reported by others also^92^. Furthermore, the hexa-histidine tag was computed to carry only a slight overall positive charge at pH 6.0^93^. The presence of D, E and K residues alongside histidine tag in the linker further nullifies positive charges and possesses a net negative charge for this segment. As observed previously^51^, our results with histidine-tag removed PrLD did not show any difference in their phase separation behavior with GRNs (Fig S11), dismissing the possibility of tag contributing to phase separation.

Earlier, we showed that coacervation with GRNs is likely driven by net charges on the protein ^51^. The charge distribution on GRNs 2, 3 and 5 shows that they are isoelectronic to negative charges, but with respect to positive charges, GRN-2 with +2 is in the middle between GRNs 3 and 5 (+4 and 0, respectively) (Fig 2), while the hydropathy of all these GRNs are remarkably similar (Table S1). This facilitated the investigations on GRN-2’s interactions with PrLD to draw biophysically meaningful inferences as GRN-3 and −5 have been shown to display a spectrum of phase transitions upon interacting with PrLD ^51^. The results presented here establish that the negative charges are the main driving forces of LLPS between GRNs and PrLD while the increase in the number of positive charges shifts the equilibrium towards aggregation (Fig 8a). To a lesser degree, hydrophobic interactions may also contribute possibly via cation -π and π - π interactions but certainly do not seem to drive LLPS. Interestingly, by simply modulating the charges on GRNs, one could seemingly control the dynamics of phase transitions and coacervation with PrLD as summarized in Figure 8b-g. When under the control of purely negative charges on GRN, PrLD undergoes LLPS to form droplets that are demixed from the solution (Fig 8b and c). This can be explained based on polyphasic linkage and ligand binding effect on phase transitions as recently postulated by Ruff and coworkers ^94^. Based on their work, preferential binding of a ligand (GRN) to the scaffold (PrLD) in dilute or dense phase can be expressed as;

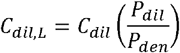

**Figure 8.**
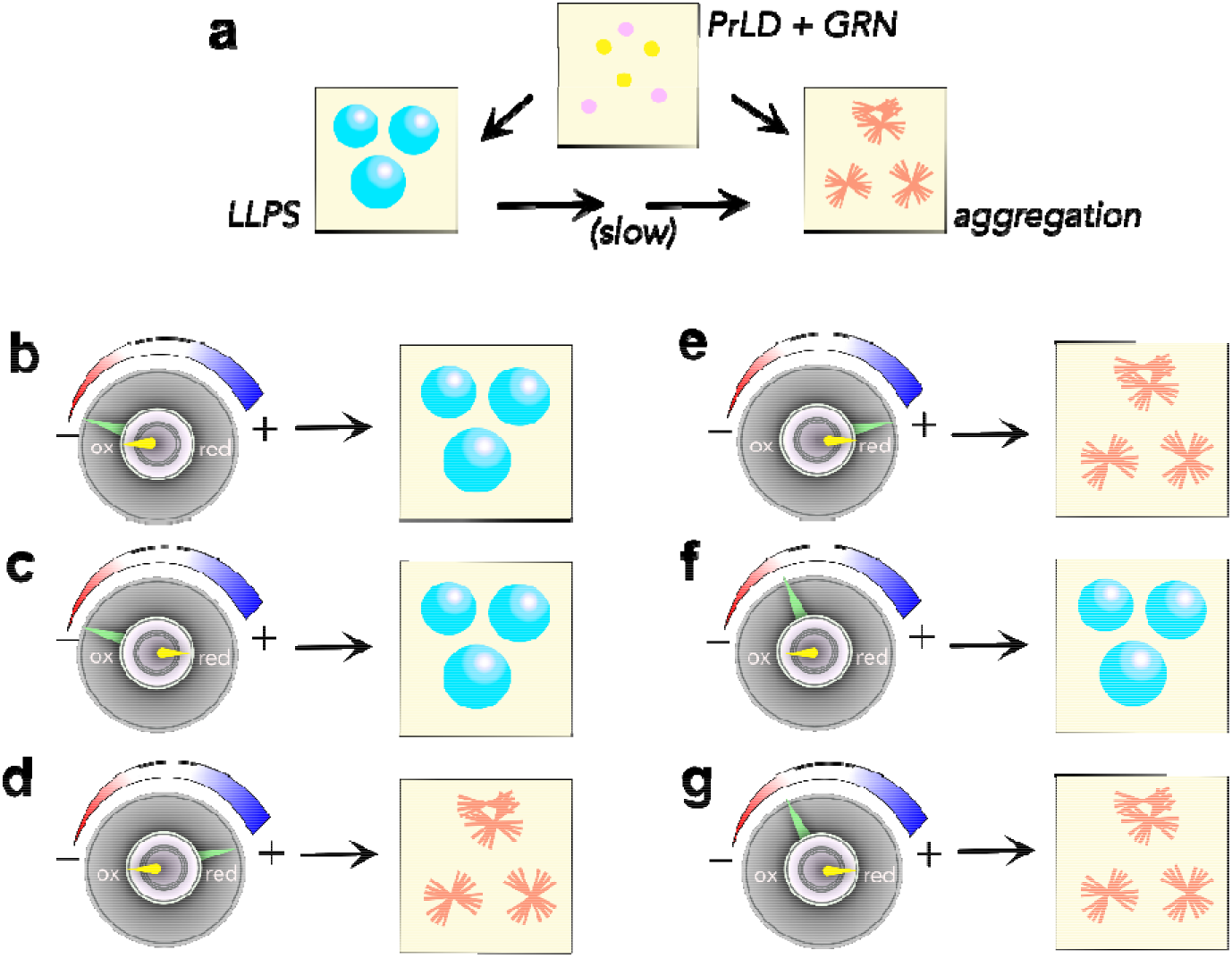
Schematic summary of coacervation between GRNs and PrLD. (a) Phase transitions of PrLD induced by GRNs. (b-g) Phase changes are coarsely controlled by electrostatic interactions (large knob) while they are finely controlled by the redox state (smaller knob) for some select conditions discovered in this report.

Where *C_dil,L_* is concentration of the scaffold in the coexisting dilute phase in ligand’s presence, *P_dil_* and *P_den_* are binding polynomials that define ligand binding to the scaffold in respective phases. If *P_dil_* > *P_den_*, i.e., binding of ligand to the scaffold is greater in the dilute phase, *C_dil,L_* > *C_den_*, LLPS gets suppressed and when *P_dil_* < *P_den_* LLPS is augmented. Therefore, negative charges potentiate multivalent electrostatic charges with PrLD to preferentially partition into the dense phase, thus promoting LLPS. Increasing the number of positive charges to four shifts the equilibrium towards the formation of solid aggregates possibly by engaging in additional salt-bridge interactions with acidic residues of PrLD, preferably in the dilute phase (Fig 8d and e). These interactions, we speculate also promote aggregation. Unfortunately, we do not have experimental evidence for this extreme case (high positive charges and no negative charges), as such a mutant posed difficulty in recombinant expression (data not shown). However, the results obtained from GRN-7 (Fig S5), which has a high number of positive charge (+8) serves as a good proxy and strengthen the validity of the idea that GRNs have modulatory capabilities especially when it is known that not all GRNs level are the same in cells ^95^.

Perhaps the most interesting aspect of this behavior is the role of cysteines and redox control. Our data indicate that when under compensatory positive and negative charge regimes in GRNs, redox conditions dictate the phase transitions (Fig 8f and g). Under fully reducing conditions, thiols in cysteines may participate in a network of hydrogen bonding interactions with Ser and/or Asn residues that are abundant in PrLD. Both these residues are conducive partners of cysteines as H-bonding donors and acceptors ^96^ and are abundant in PrLD (together accounting for ~25% of the sequence). Empirically, it is evident that these additional valences increase solute-solute interactions that shift the equilibrium out of the liquid-like state towards an aggregated state. The possibility of such a scenario is supported by the behavior of the alkylated forms of GRN-2 (Fig 5), where the polar acetamide adducts are capable of engaging in similar interactions. But under oxidizing conditions, the disulfide bonded cysteines preclude such interactions are abrogated, and therefore maintain weak and transient interactions that promote LLPS and a demixed state. From this, we surmise that the positively charged residues and thiols help fine-tune the phase transitions. It is also noteworthy that not an abrupt, but a continuum of phase changes is observed when traversing between one end of the redox spectrum to the other (Fig 5), which indicates that even under partially reducing conditions, LLPS can occur to some degree. Therefore, it seems plausible that GRN-PrLD droplets and aggregates could co-exist in the cytoplasm which largely presents a reducing environment.

The data presented here unambiguously establishes coacervation between PrLD and GRNs, yet, the absence of GRNs in SGs within PrLD or full-length TDP-43 expressing neuroblastoma cells seems to counter-intuitive at the outset. However, as observed by the ternary interactions between PrLD, GRN-2 and RNA, the reason for the inability of GRNs to partition into SGs is clear. GRN within the condensates of PrLD simply gets displaced by a stronger electrostatic ligand, RNA, which abrogates the possibility of the three co-existing within the SGs. Given that SGs also contain many other phase separating proteins such as TIA1 ^57^, G3BP1 ^97^, hnRNPA1 ^98^ etc., the likelihood of GRNs partitioning into SGs is low. Furthermore, it is also interesting to observe that RNA is able to out-compete GRN-2 in reducing conditions where it forms insoluble aggregates with PrLD (Fig 7). This, along with the observed colocalization of GRN and PrLD as puncta in both non-stress and stress conditions, suggest that SGs could partly mitigate the potential toxicity posed by GRN-PrLD inclusions, which eventually transition to amyloids, with an intervening gelated phase.

This report brings out the first evidence for the cellular uptake of extracellular GRNs and their cytoplasmic colocalization with PrLD. The exquisite control and tuning of coacervation of PrLD by GRNs via electrostatics and redox flux present intriguing possibilities by which GRNs can influence pathology. During acute inflammation, when GRNs are known to be generated in abundance by microglia and astrocytes ^42, 99–101^, these pro-inflammatory molecules are taken up by neurons where they modulate the dynamics of cytotoxic proteolytic fragments of TDP-43. Even with haploinsufficiency of PGRN, which is a risk factor for FTLD, it can be conjectured that inflammation could lead to augmented production of GRNs that could compensate for the loss in PGRN ^42^. Perhaps the most interesting aspect when observing the sequence of the seven GRNs is that they seem to have a perfect combination and spectrum of positive and negative charges to eloquently modulate TDP-43 phase transitions. It also seems to be that the extracellular cleavage of PGRN may be asymmetric leading to a greater abundance of one GRN over another^95^, which could play in the modulation of TDP-43. Nevertheless, precisely what cellular consequences do these inclusions and coacervates possess remain unclear at this time. A few possibilities include induction of mitochondrial, lysosomal, and autophagic dysfunction. Some of these will be unearthed in the coming years with the establishment of complex interactions and phase transitions between GRNs and TDP-43 PrLD as described in this report.

## Materials and Methods

### Recombinant expression and purification of proteins

#### GRNs

GRNs (GRN-2, GRN-2(+/-), GRN-2(C/A), GRN-3, and GRN-5) were recombinantly expressed and purified as previously described, where GRN-3 was expressed in *Escherichia coli* SHuffle cells (New England Biolabs) while other GRNs were expressed in Origami 2 DE3 cells (Invitrogen) ^51^. Briefly, GRNs were expressed as fusion constructs containing an N-terminal thioredoxin-A, and His_6_ tag with an intervening thrombin cleavage site. The fusion constructs were purified using immobilized-nickel affinity chromatography. The purified construct was cleaved using restriction grade bovine thrombin (BioPharm Laboratories) at 3 units/1 mg of protein for 24 h at room temperature to separate the fused tags. For GRN-2ΔAro, purification protocol was suitably modified to minimize thiol oxidation; briefly, buffers were degassed prior to use and the purification was carried out at room temperature to reduce oxygen solubility. The sample was fractionated using semipreparative Jupiter 5 μm-10 x 250 mm C18 reverse-phase HPLC column (Phenomenex) using a gradient elution of 60%–80% acetonitrile containing 0.1% TFA. Fractionated protein was lyophilized and stored at −20°C. For the generation of 2-alkylated GRN-2 i.e. GRN-2 with two-free thiols that are alkylated, the protein was incubated with 10 molar excess of iodoacetamide post-thrombin cleavage for a period of 24 h at room temperature. Protein was then fractionated using reverse-phase HPLC as before. For the generation of 12-alkylated GRN-2 i.e. GRN-2 with all twelve thiols alkylated, the protein was incubated with 12x molar excess of TCEP post-thrombin cleavage for a period of 12 h at room temperature followed by incubation with 10 molar excess of iodoacetamide for 24 h at room temperature. Protein was then fractionated using reverse-phase HPLC as before. The purity of protein was confirmed using matrix assisted laser desorption-ionization time-of-flight mass spectroscopy (MALDI-ToF MS).

#### TDP-43 PrLD

PrLD was expressed and purified as described previously ^51^. The plasmid for TDP-43 PrLD was a gift from Dr. Nicolas Fawzi at Brown University (Addgene plasmid 98669, RRID: Addgene 98669). Briefly, the protein was expressed as a fusion construct with an N-terminal His_6_ tag followed by tobacco etch virus (TEV) protease cleavage site in *E.coli* BL21 DE3 Star cells (Invitrogen). The fusion construct was purified using immobilized-nickel affinity chromatography. Purified protein was concentrated using Amicon Ultra-Centrifugal units (Millipore) and flash frozen for storage at −80°C or used immediately for experiments.

### Preparation of proteins and RNA

#### GRNs

Lyophilized protein was resuspended in required buffer (20 mM MES, pH 6.0 or 20 mM ammonium formate, pH 4.5) and the concentration was estimated spectrophotometrically at 280 nm with extinction coefficients of 6250 M^-1^ cm^-1^ for GRN-2, GRN-2(+/-) and GRN-3, 5500 M^-1^ cm^-1^ for GRN-2(C/A), 7740 M^-1^ cm^-1^ for GRN-5 or at 215 nm for GRN-2ΔAro with an extinction coefficient of 90000 M^-1^ cm^-1^. The number of free thiols within GRNs was estimated using Ellman’s assay and by alkylation with iodoacetamide, as described previously^102^. The reduced forms of GRNs were generated by incubating the freshly purified proteins with 12x molar excess of tris(2-carboxyethyl)phosphine (TCEP) at room temperature for 2-4 h or at 4°C for ~12 h. Fluorescent labeling of GRNs was performed using HiLyte™ Fluor 405 succinimidyl ester (Anaspec) or HiLyte™ Fluor 647 succinimidyl ester for FRAP studies on GRNs. Proteins were incubated with 3 molar excess of dyes at 4°C for ~12 h and excess dye was excluded using clarion MINI Spin Columns, Desalt S-25 (Sorbent Technologies Inc).

#### PrLD

Before the experiments, the protein was buffer-exchanged into 20 mM MES, pH 6.0 or 20 mM ammonium formate, pH 4.5, using PD SpinTrap G-25 desalting columns (Cytiva) and the concentration was estimated using an extinction coefficient of 19480 M^-1^ cm^-1^ at 280 nm. Fluorescent labeling of PrLD was performed using HiLyte™ Fluor 647 succinimidyl ester (Anaspec) using a similar protocol as described above for GRNs.

#### RNA

Lyophilized poly-A (Sigma Aldrich) was resuspended in deionized, sterilized water at a concentration of 1 mg/mL. The concentration of the stock was estimated by considering a value of 1 absorbance unit to correspond to 40 μg of RNA. The prepared stock was flash frozen and stored at −80°C. Aliquots of prepared stock were thawed and used immediately for experiments.

### Cell growth, transfection and colocalization analysis

SH-SY5Y neuroblastoma cells (ATCC, Manassas, VA) were grown in DMEM:F12 (1:1) media containing 10% FBS (Gibco, Thermo Scientific) and were maintained in humidified condition at 37 °C with 5.5% CO_2_. Cells were seeded 24 hours before transfection. Cells were transfected with PrLD-SBFP2 or wtTDP43tdTomato plasmid using the TransIT-X2^®^ dynamic delivery system, Mirius (1:3) in Opti-MEM media (Thermo scientific). Cell confluency was allowed to reach 70-80% prior to transfection. After 24 h, cells were gently washed for two times with fresh media and 500 nM of fluorescently labeled recombinant GRNs were added. Cells were incubated with GRN-containing media for 24 hours. Following this, media was replaced with GRN-devoid media and cells were incubated for a further 1 h. For inducing stress, sodium arsenite was added at a final concentration of 0.5 mM and incubated for 30 minutes. Cells were stained with nuclear (NucSpot^®^ Live 650, Biotium) or lysosomal (Lysoview™ 650, Biotium) markers prior to imaging at 40X magnification using Leica STELLARIS-DMI8 microscope. For colocalization analysis of GRNs and PrLD, the respective channels for PrLD and GRNs were converted to 16-bit grayscale format and region of interest was selected. Images were processed using Coloc2 parameter in FIJI imageJ software using Costes threshold regression, and Manders’ tM1 values were reported. All the confocal images were processed using Adobe illustrator and data were processed using OriginPro 8.5 software.

### Immunofluorescence

For immunofluorescence experiment, cells were plated in a 12 well plate 24 h prior to transfection with PrLD-SBFP2 as described above. After incubating with fluorophore labeled-GRNs for 24 h, media was replaced, and experiments were carried out in both stress and non-stress conditions. For induction of stress, 0.5 mM sodium arsenite was added to the media and incubated for 30 minutes followed by washing with 1x PBS. Cells were fixed in 4% paraformaldehyde for 20 minutes and washed twice with 1x PBS. These were further permeabilized with 0.2% Triton X-100 in 1x PBS for 20 minutes. Cells were washed and blocked using 3% BSA for 2 h. Cells were incubated with anti-TIA1 primary antibodies (Cell Signaling Technology, TIAR XP^®^ Rabbit mAb #8509) and anti-rabbit secondary antibodies (Cell Signaling Technology, Alexa Flour^®^ 488 conjugate #4412) and imaged following the addition of antifade mounting media at 40X magnification on Leica STELLARIS-DMI8 microscope.

### DIC Microscopy and FRAP analysis

DIC and fluorescence microscopy images were acquired on Leica STELLARIS-DMI8 microscope. The assays were performed in an optical bottom 96 well-plate (Thermo) and were covered with an optically clear sealing tape (Nunc, Thermo Scientific) to prevent evaporation. The reactions were initiated at room temperature and were visualized within 10-15 mins of mixing. Brightfield and fluorescence images were acquired at a magnification of 40x or 63x with an oil immersion objective. For microscopically monitoring the ternary interactions between GRN-2, PrLD and RNA; we initially generated heterotypic condensates of GRN-2 and PrLD, followed by titration with desired volumes of the RNA stock. Images were acquired after an equilibration time of ~10-15 mins after each titration. The RNA stocks were prepared at a high initial concentration to minimize dilution effects upon titration. For fluorescence imaging and FRAP studies, samples were prepared by mixing the required concentration of proteins in an optical-bottom 96 well plate in the presence of 1% fluorophore labelled proteins. For FRAP studies, fluorescence intensities were acquired pre- (~2 seconds) and post-bleach (~30 seconds) at an interval of 68 μs. Bleaching period was varied from 10-20 seconds with a laser intensity of 100% while the laser intensity for imaging was set at 2-10%. The fluorescence recovery data was processed using Leica LasX program and Origin 8.5 graphing software. The recovery curves of at least three independent and individual samples were averaged and normalized with respect to the pre-bleach fluorescence intensities. To discern the rate constants of recovery kinetics post-bleach the initial-rate method was utilized ^103–104^. Briefly, period of fluorescence recovery 1 second post-bleach was evaluated via a linear fit with the slope of the line providing a rate constant, *k* (s^-1^). Rate constants were obtained by evaluating three FRAP recovery curves for each sample. For time-course experiments, FRAP analysis was performed on samples at initial time (0 h) and after incubation of 24 h at 37°C under quiescent conditions. A noteworthy point is that we observed significant photobleaching of HiLyte-405 when performing FRAP analysis of GRNs labeled with the fluorophore precluding the use of this dye in this study. To obtain this data, we then cross-labeled GRNs with HiLyte-647 (for performing FRAP studies) and PrLD with HiLyte-405 (only for visualization).

### Post imaging analysis

#### Droplet area distribution

Area of the droplets formed within various samples was ascertained using the ImageJ program. Briefly, the fluorescence images acquired for the respective samples were subjected to color thresholding at default values in the program. Area of droplets within the threshold images were then extracted using the included particle analyzer tool of the program. A lower limit of either 50 or 200 (pixel^2^) (depending on image magnification) was placed on area of particles to be analyzed so as to exclude detection of small artifacts. Areas obtained in pixel^2^ were converted into μm^2^ by considering the scale-bar. The obtained data was then plot as a histogram overlayed with normal distribution curve using Origin 8.5. A set of 100 droplets were enumerated from at least two different fields for each sample.

#### Fluorescence intensity profiles

To qualitatively ascertain the displacement of GRN-2 from condensates upon titration with RNA, we determined the fluorescence intensity profiles of the fluorophore labeled proteins from the images of the respective samples using the ImageJ program. Intensity profiles were obtained by mapping a line segment across the width of a droplet and extracting the intensities of the fluorophores (HiLyte 405 for GRN-2 and HiLyte 647 for PrLD) using the plot-profile tool. The plotprofile tool provides intensity values within an arbitrary window of 0-80 units, with highly fluorescent samples being cut-off at the upper limit. The fluorophore intensities obtained across a pixel scale were then converted into μm using the scale-bar. The fluorescence intensity profile was extracted from individual droplets for each sample but are representative of the entire field containing at least 20 different such droplets.

### Turbidimetric assays

Turbidity measurements were performed on a BioTek Synergy H1 microplate reader. Reactions were initiated and allowed to equilibrate at room temperature for 3-10 mins before each measurement. For titration experiments with GRNs, NaCl, or RNA, stocks were prepared at high concentrations to minimize dilution of samples. For establishment of phase diagrams, a boundary value of 0.140 OD_600_ was set and readings above it were considered to undergo phase transition. Temperature dependent phase diagram was established by equilibrating samples at respective temperature (4°C, room temperature) for 15 mins, while temperatures above 30°C were achieved using the internal temperature control capability of the plate reader. Data processing was performed on Origin 8.5. At least independent data sets (n=3) were averaged.

### Aggregation assay

To monitor the amyloid-formation of PrLD in presence of GRN-2 or rGRN-2, thioflavin-T (ThT) fluorescence assays were performed on BioTek Synergy H1 microplate reader. The aggregation of PrLD was monitored in the presence of different molar concentrations of GRN-2 or rGRN-2 in separate reactions in the presence of 15 μM ThT dye. The reactions were monitored for a period of 24 h under quiescent conditions at 37°C and are representative of three independent data sets.

### MALDI-ToF MS

For confirming the purity and for characterization of proteins used, MALDI-ToF MS spectroscopy was performed on a Bruker Daltonics Microflex LT/SH TOF-MS system. Prepared samples were spotted onto a Bruker MSP 96 MicroScout Target microchip with a 1:1 ratio of sample:sinapinic acid matrix in saturated acetonitrile and water. Instrument calibration was performed using Bruker Protein Calibration Standard I (Bruker Daltonics). Alkylation assays of GRNs using iodoacetamide were performed as described ^105^.

### In-silico analysis and figure preparation

Redox-dependent disorder scores for individual proteins were obtained using IUPred2A platform^76^. Specifically, the FASTA sequence of each protein was evaluated by coupling the IUPred2 long disorder algorithm with a Redox-state tool ^77^. The hydropathy indices for GRNs were obtained using the CIDER platform ^106^. The obtained figures were processed using the Adobe Illustrator CC suite and Affinity Designer^®^.

## Acknowledgements

The authors would like to thank the following agencies for financial support: National Institute of Aging (1R56AG062292-01) and the National Science Foundation (NSF CBET 1802793) to VR. The authors also thank the National Center for Research Resources (5P20RR01647-11) and the National Institute of General Medical Sciences (8 P20 GM103476-11) from the National Institutes of Health for funding through INBRE for the use of their core facilities. The authors would also like to thank the INBRE facility manager, Dr. Jonathan Lindner for his gracious help in using the confocal microscope.

## Author contributions

VR conceptualized the project; AAB performed protein expression and biophysical experiments, SD performed the cell culture experiments; VR, AAB and SD participated in intellectual discussions and manuscript writing and editing.

## Competing interests statement

The authors declare that they have no financial or non-financial interests.

## TOC figure

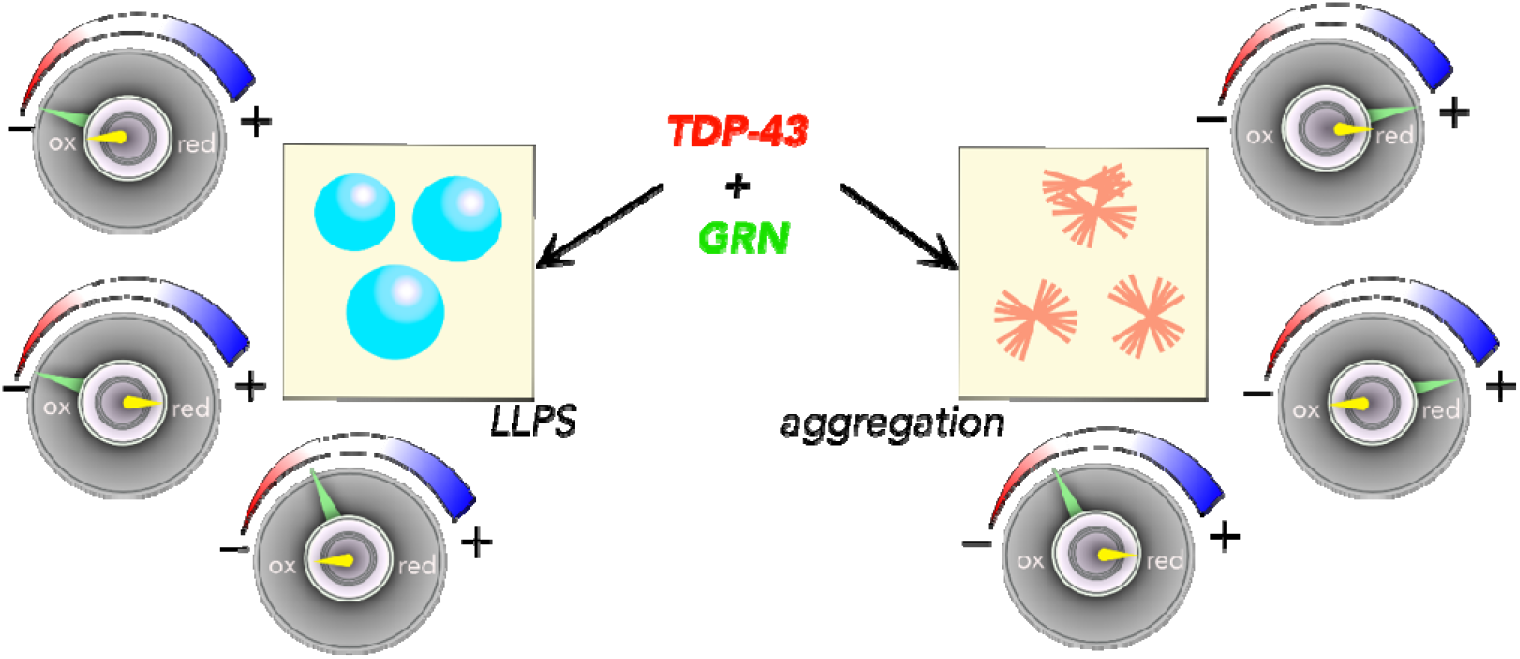

## SUPPLEMENTARY FIGURES

**Figure S1.**
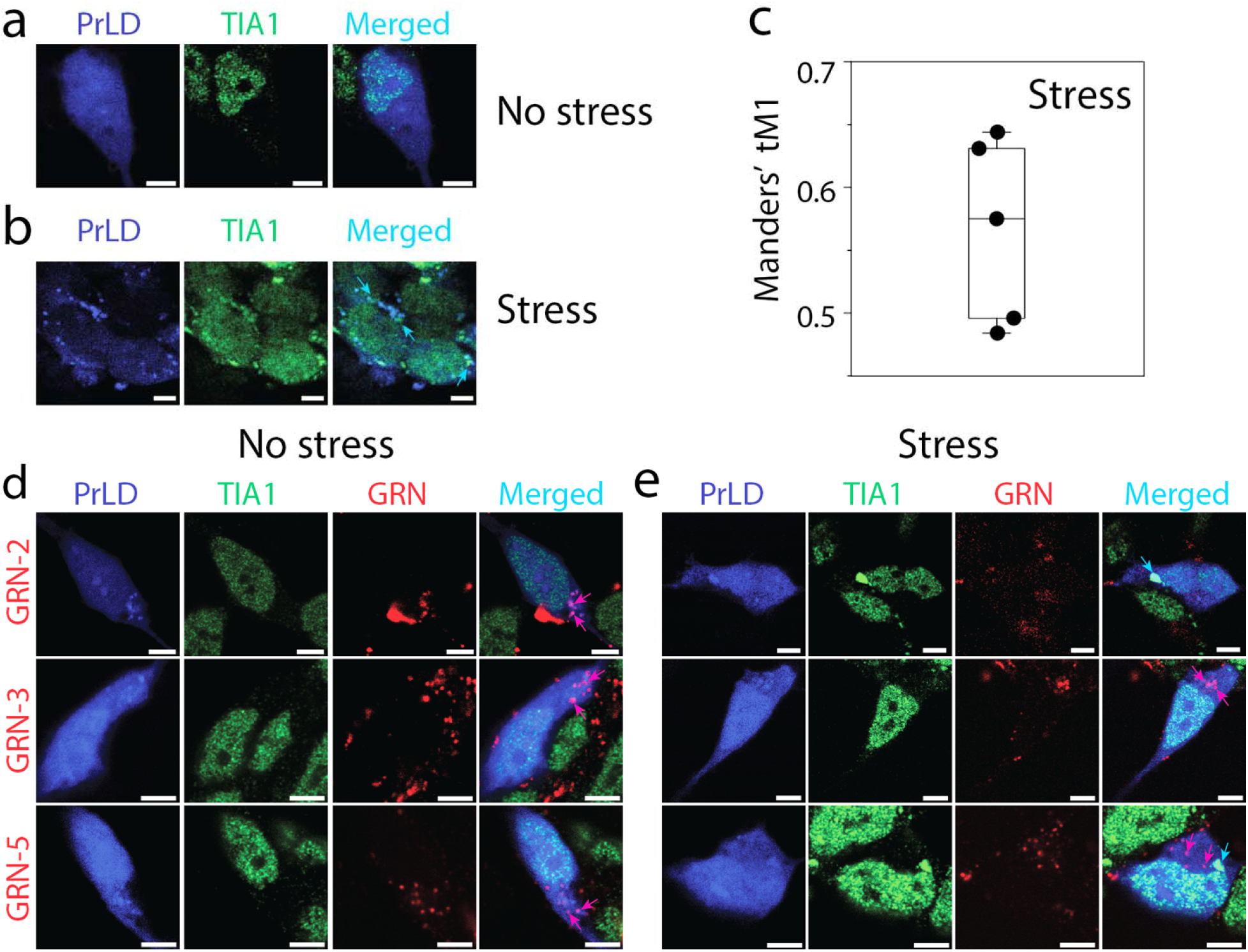
Localization of GRNs and PrLD within SGs. a-b) Immunofluorescence imaging of PrLD along with GRNs using TIA-1 antibodies with and without stress. PrLD-SBFP2 colocalization study with TIA1 in non-stressed cells (a) and sodium arsenite-induced stressed cells (b) (cyan arrows). C) Manders tM1 derived from five independently observed cells. d-e) Colocalization analysis of GRNs labeled with Hilyte-647, PrLD and/or TIA-1 without stress (d) and with stress (e). Cyan colored arrows indicate the colocalization of PrLD and TIA1, while pink colored arrows represent colocalization of PrLD and GRNs. A three-way colocalization between GRN, PrLD and TIA1 was not observed. Scale bar represents 5 μm.

**Figure S2.**
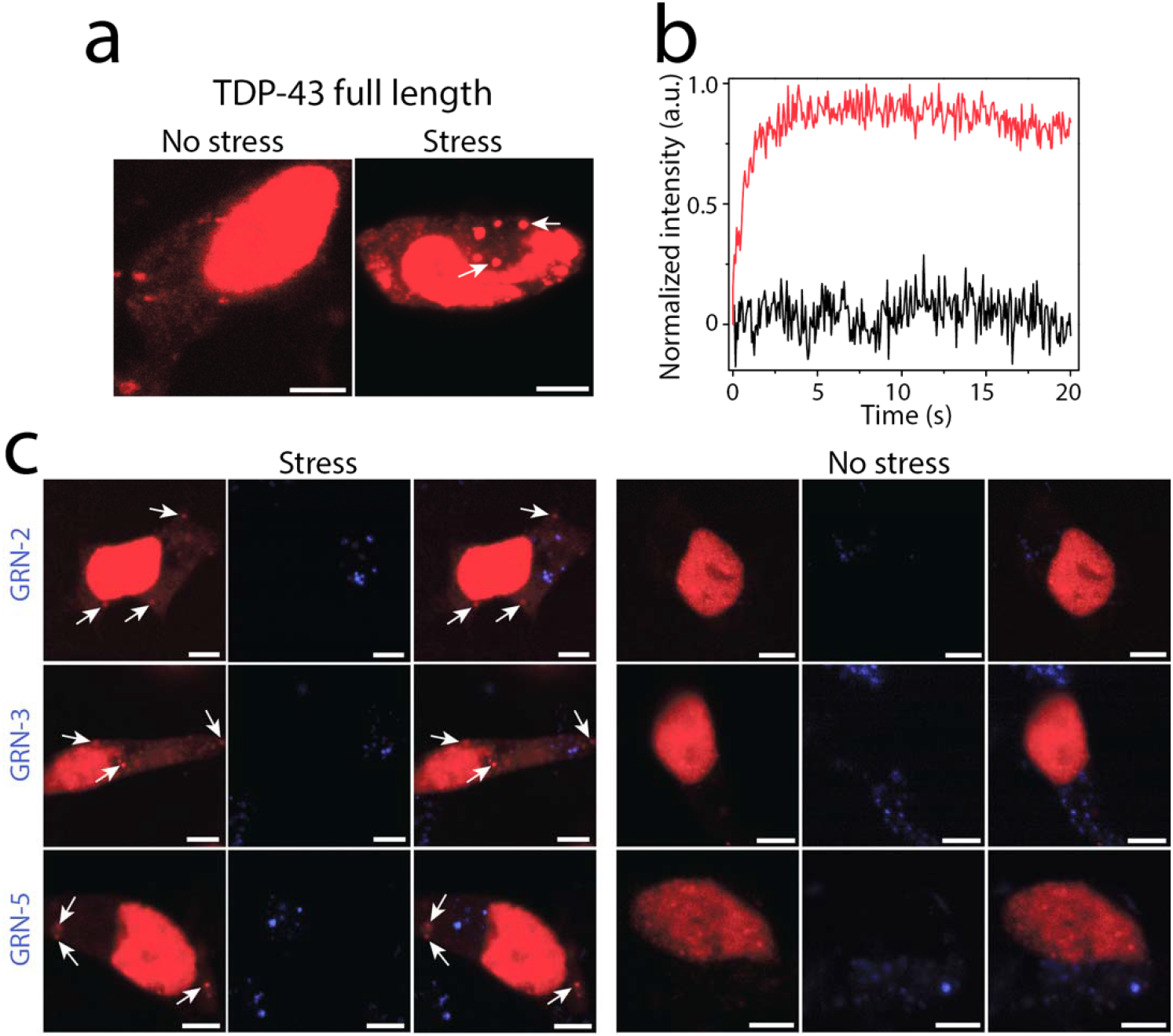
Live cell analysis of SGs formed by full-length TDP-43 in SH-SY5Y cells. Wild-type TDP43 tdTomato with and without stress in SH-SY5Y cells (a). White arrows indicate the SGs formation in sodium arsenite treated cells. Normalized FRAP data of TDP-43 SGs and solid-like aggregates in sodium arsenite treated cells 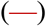 and untreated cells (—) respectively. c-d) Confocal images of wtTDP43tdTomato transfected cells in presence of GRNs without (c) and with stress (d). Arrows in stressed cells indicate SGs containing TDP-43. No colocalization between GRNs and TDP-43 was observed. Scale bar represents 5 μm.

**Figure S3.**
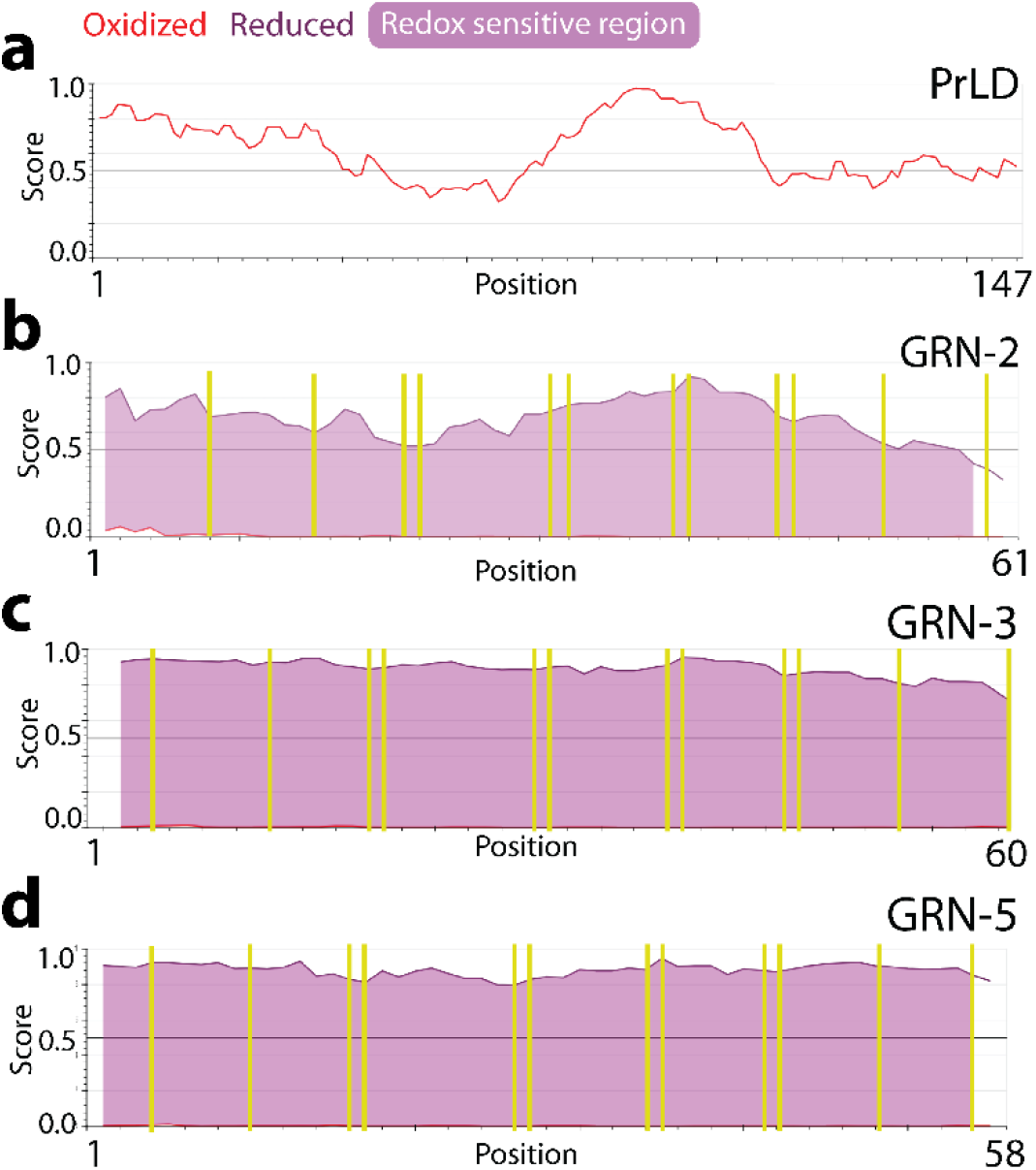
Redox-controlled disorder propensities of PrLD and GRNs. Structural disorder score of PrLD and GRNs estimated in a redox-dependent context using IUPred2A. Redox sensitive regions (shaded, purple) are calculated based on differences in disorder scores of the oxidized (red) and reduced (dark purple) forms of the proteins. Calculated scores above the 0.5 threshold signify disorder within the region. Cysteine residues are marked within the sequence of GRNs (yellow bars).

**Figure S4.**
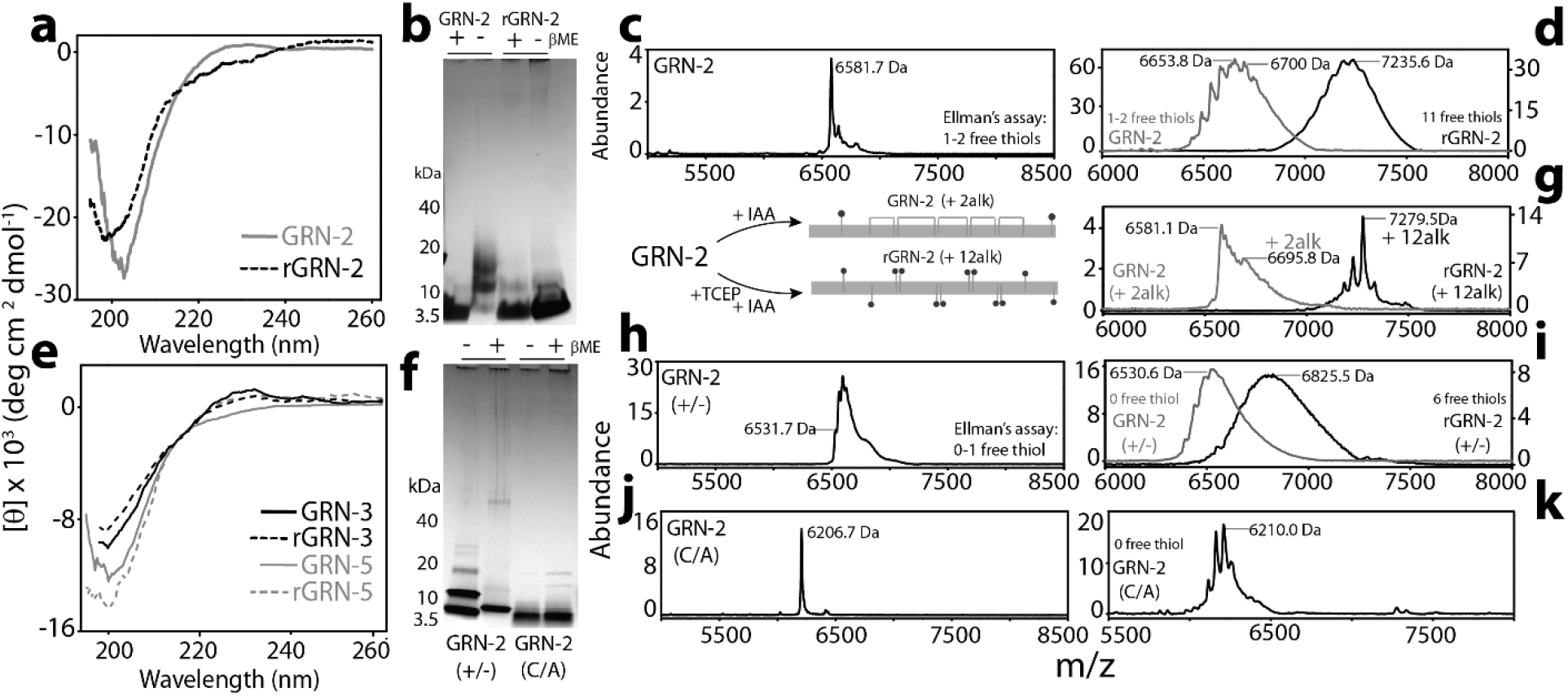
Biophysical characterization of GRNs. a) Far-UV circular dichroism (CD) spectra of GRN-2 and rGRN-2 displaying a characteristic profile of a structure dominated by random coils. b) SDS-PAGE analysis of GRN-2 and rGRN-2 the presence or absence of β-mercaptoethanol (βME) performed under denaturing conditions. c) MALDI-ToF MS graph of GRN-2 shows a peak at 6581.7 Da which corresponds to the monomeric form of the protein (theoretical mass: 6590.5 Da). Ellman’s assay performed on the fraction reveals the presence of 1-2 free thiols. d) Free thiols in GRN-2 and rGRN-2 determined by alkylation with iodoacetamide and analyzed using MALDI-ToF MS. Alkylation of a free thiol by iodoacetamide leads to the addition of a thioether adduct (+ 59.0 Da). e) Far-UV CD spectra of the redox forms of GRN-3 and GRN-5. f) SDS-PAGE analysis of the mutant forms of GRN-2; GRN-2(+/-) and GRN-2(C/A) in the presence or absence of βME under denaturing conditions. g) Alkylated forms of GRN-2 were generated by either capping the two free-thiols in the oxidized form (GRN-2 2alk) or all twelve thiols in the reduced form (GRN-2 12alk) using iodoacetamide and were subsequently subjected to MALDI-ToF MS for analysis. h) Characterization of GRN-2(+/-) using MALDI-ToF MS shows a peak at 6531.7 Da corresponding to monomeric form of the protein (theoretical mass: 6536.4 Da). Two additional peaks were observed beside this corresponding to two, highly oxidized sulfur atoms in the cysteine (Sulfur-dioxide; +32 Da). Ellman’s assay reveals the presence of about 1-free thiol in the protein i) Estimation of free thiols in the oxidized and reduced form of GRN-2(+/-) using alkylation assay. j) Characterization of GRN-2(C/A) using MALDI-ToF MS shows a peak at 6206.7 Da corresponding to monomeric form of the protein (theoretical mass: 6205.8 Da). k) Alkylation assay performed on GRN-2(+/-) analyzed using MALDI-ToF MS.

**Figure S5.**
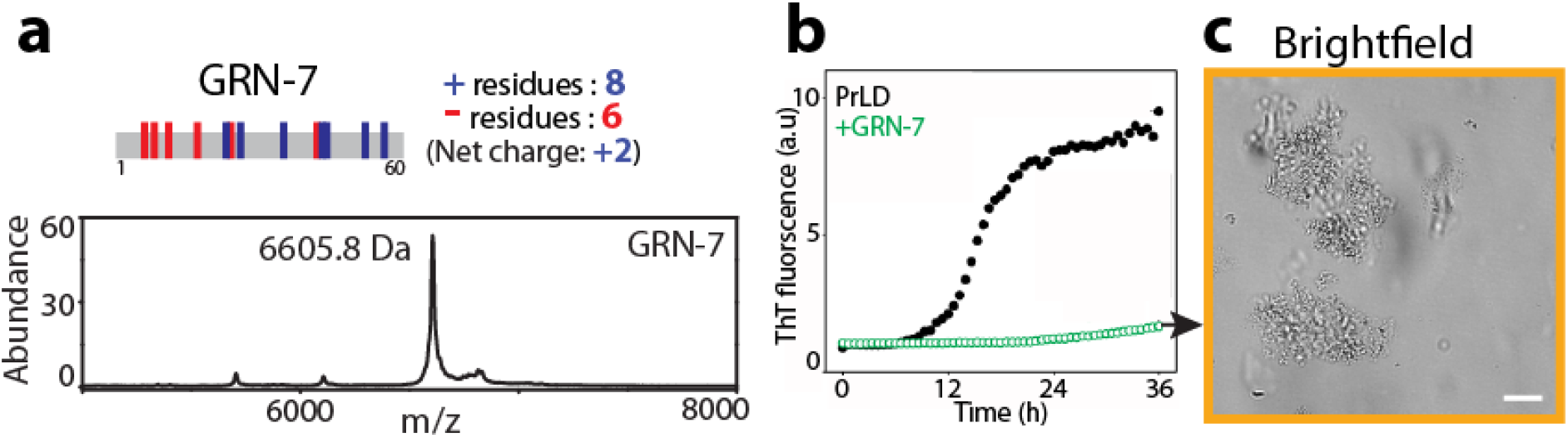
Interaction of GRN-7 with PrLD. a) Sequence of GRN-7 annotated with acidic and basic residues and the net charge at neutral pH. MALDI-ToF MS spectrum of oxidized GRN-7 shows a peak at 6605.8 Da corresponding to the monomeric protein (theoretical mass: 6615.4 Da). b) The effect of 40 μM GRN-7 on the aggregation of 20 μM PrLD was monitored in the presence of 15 μM ThT for a period of 36 h at 37°C under quiescent conditions. c) DIC micrograph of the sample containing GRN-7 and PrLD from b) showing the presence of solid-aggregates (liquid-solid phase separation, saffron box). Scale bar represents 20 μm.

**Figure S6.**
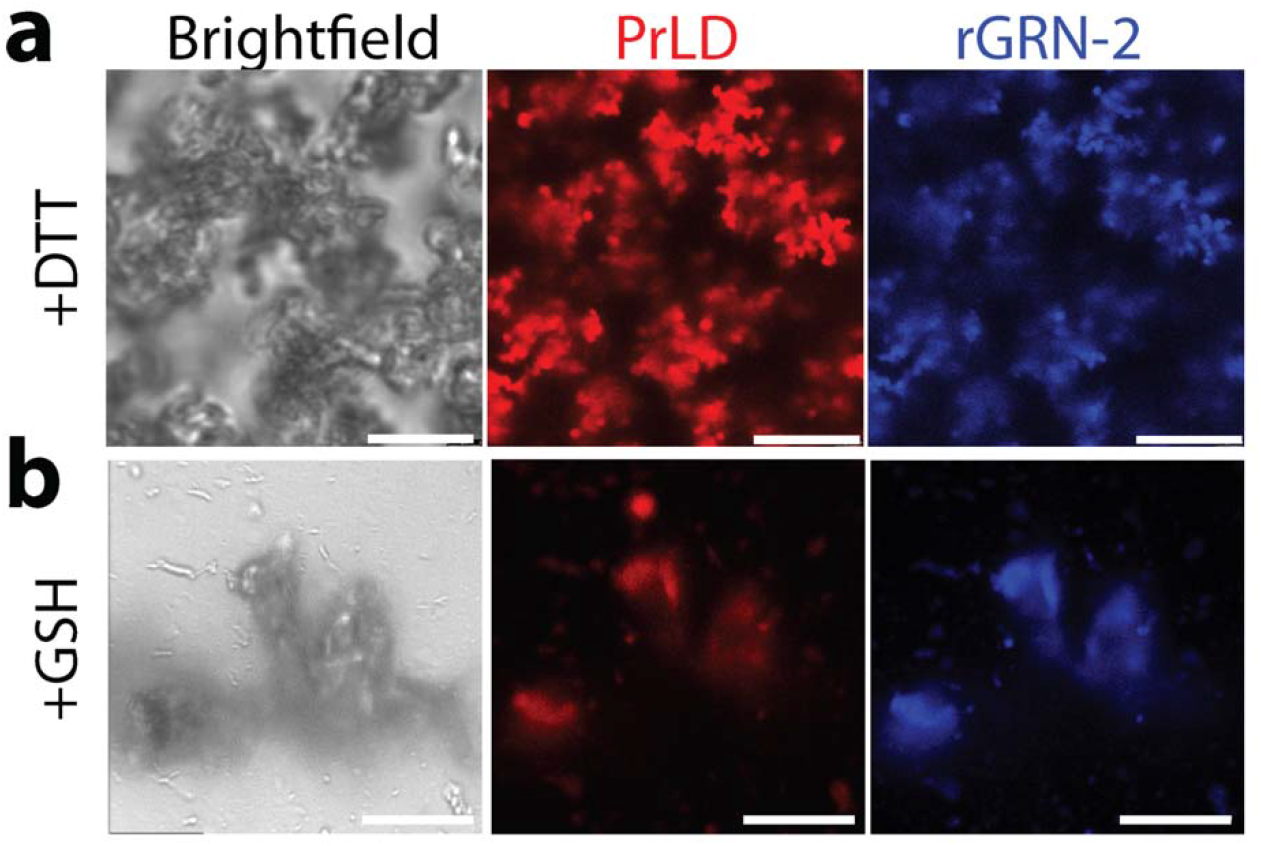
Reduction of GRN-2 with alternative reducing agents. a-b) Confocal micrographs of mixtures containing 20 μM PrLD with 40 μM rGRN-2 reduced using 12 molar excess of dithiothreitol (DTT) (a) or gluthathione (GSH) (b). Reactions contain 1% fluorophore labeled proteins for microscopic visualization. Reactions were initiated at room temperature and imaged within 15 minutes. (n = 3).

**Figure S7.**
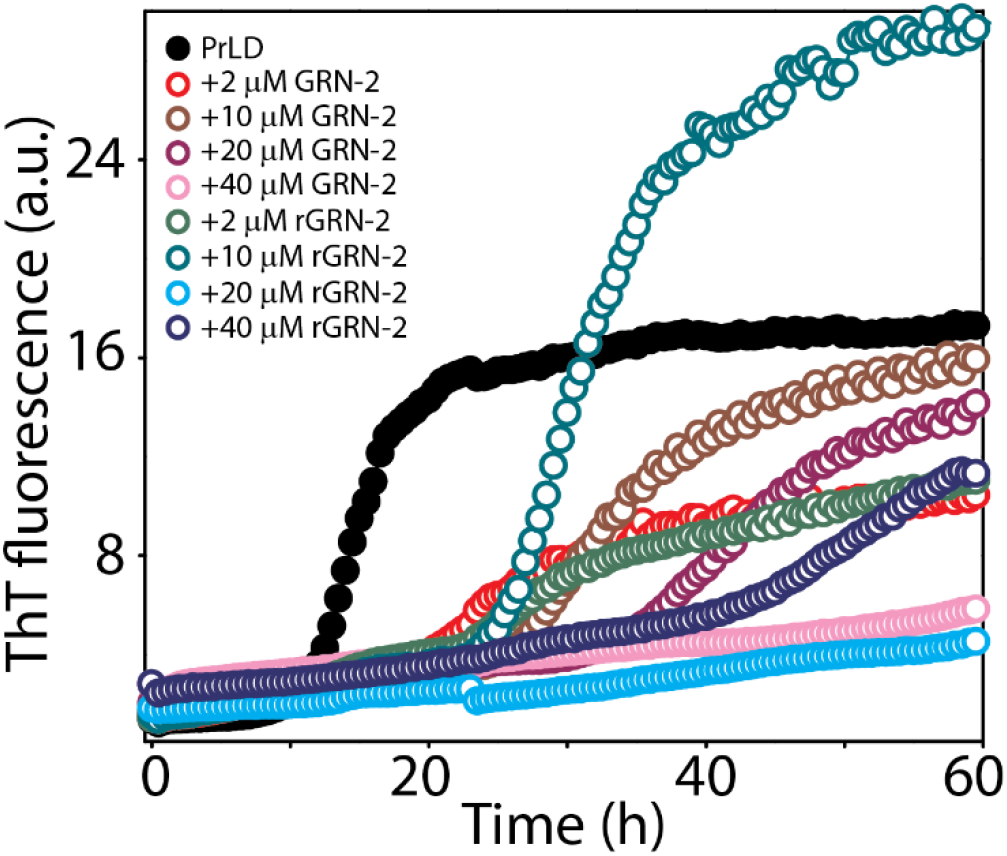
Formation of ThT-positive species of PrLD in presence of GRN-2. The amyloid formation of 20 μM TDP-43 PrLD alone (●) or in presence of varying concentrations (2-40 μM) of GRN-2 or rGRN-2, buffered in 20 mM MES, pH 6.0, monitored using 15 μM ThT for a period of 60 h at 37 °C under quiescent conditions.

**Figure S8.**
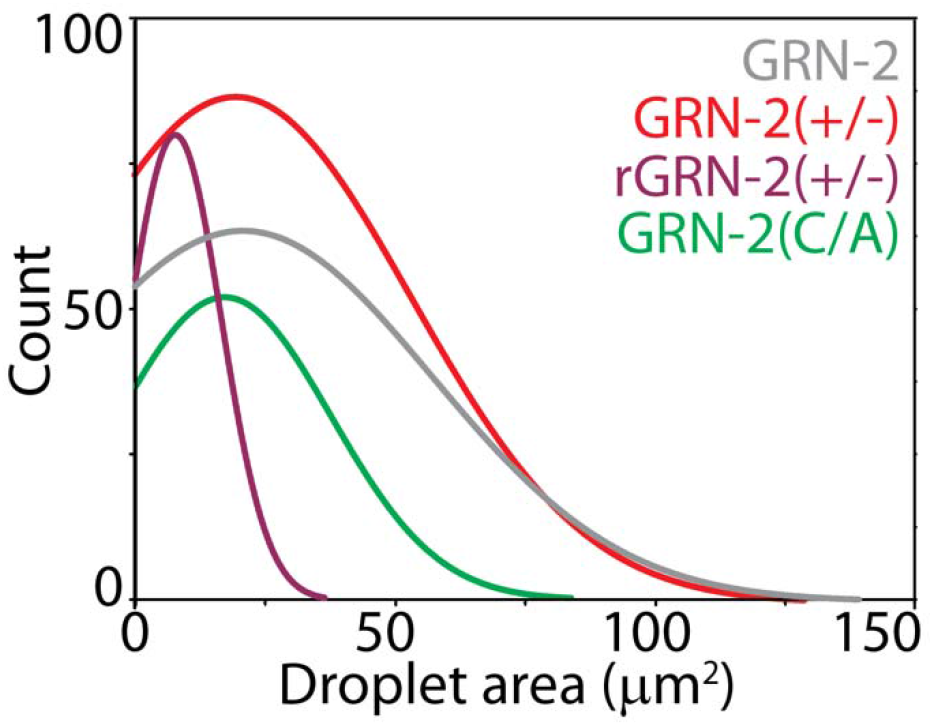
Comparison of the droplet area distributions of GRN-2 and its mutants. Micrographs of reactions depicted in Fig 5 were subjected to droplet area distribution analysis as described previously using the imageJ platform. Samples of 20 μM PrLD with 40 μM GRN-2 (gray), GRN-2(+/-) (red), rGRN-2(+/-) (purple) and GRN-2(C/A) (green) were considered for analysis with a minimum of 100 droplets enumerated. The normal distribution curves for each sample depicted were generated using Origin 8.5 graphing software.

**Figure S9.**
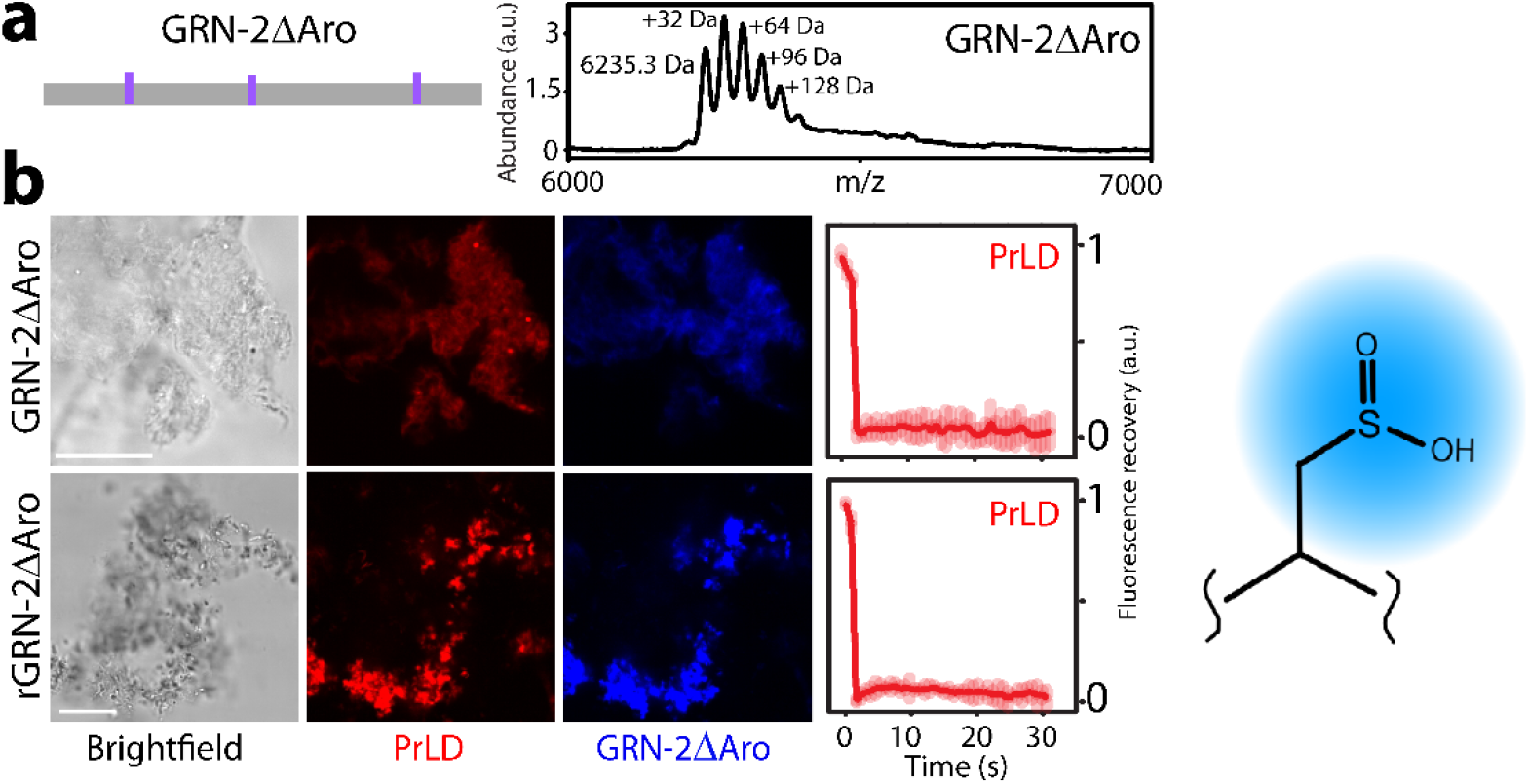
Interaction of GRN-2ΔAro mutant and PrLD. a) Sequence of GRN-2ΔAro annotated with alanine residues which replaced aromatic amino acids (Trp, Tyr and Phe) in the native sequence. MALDI-ToF MS spectra of purified protein (M.W. 6247 Da) indicating the presence of multiple sulfonated species. b) Micrographs of reactions containing 20 μM PrLD with 40 μM GRN-2ΔAro or rGRN-2ΔAro. The samples were generated by mixing 1% (molar) fluorophore-labeled proteins with unlabeled proteins buffered in 20 mM MES, pH 6.0. The samples were analyzed using FRAP analysis and resultant curves were normalized with respect to pre-bleached fluorescence intensity. Scale bar represents 20 μm. Schematic depicting the hydrophilic sulfonated thiol.

**Figure S10.**
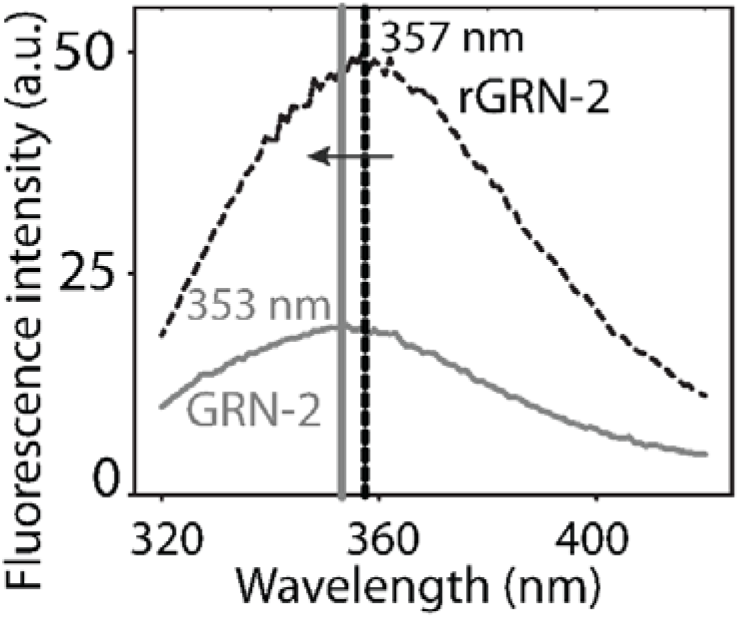
Tertiary structure analysis of GRN-2. Intrinsic tryptophan fluorescence of 20 μM GRN-2 (gray) and rGRN-2 (black) indicated a blue-shift of ~4 nm in the emission maxima of GRN-2 (353 nm, solid gray line) compared to rGRN-2 (357 nm, dashed black line).

**Figure S11.**
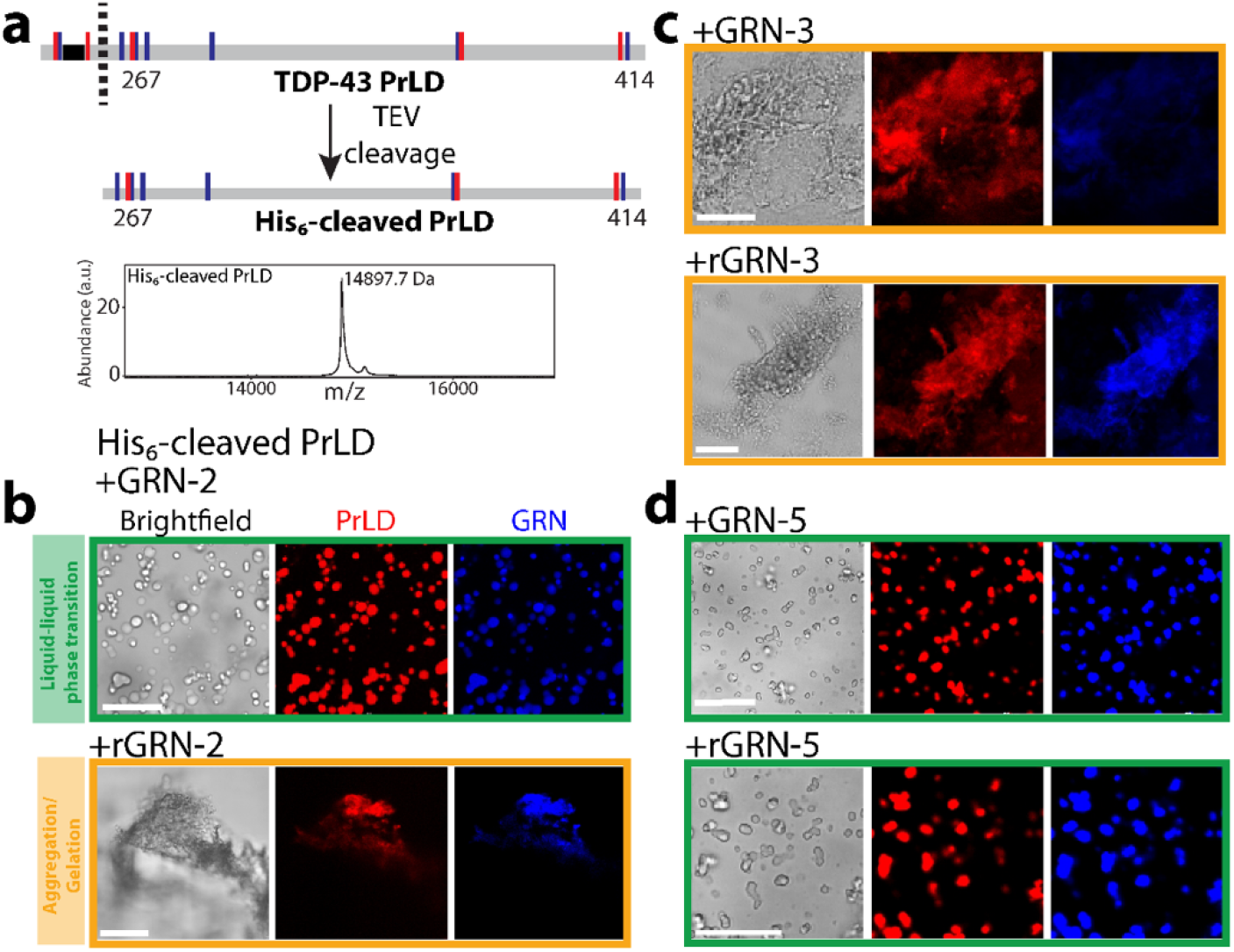
Interaction of GRNs with His_6_ cleaved PrLD. a) PrLD devoid of His_6_ tag was generated by the use of TEV protease. MALDI-ToF MS spectra of His_6_-cleaved PrLD (M.W. 14886 Da) displaying a peak at 14897.7 Da. b-d) Fluorescence micrographs of mixtures containing 20 μM His_6_-cleaved PrLD and 40 μM GRN-2 (b), GRN-3 (c) or GRN-5 (d). The green box indicates liquid-liquid phase transition and saffron box indicates aggregation or gelation. Scale bar represents 20 μm.

**Table S1:** Electrostatic and hydrophobic character of proteins under study. Listed here are the net charges and net charge per residue on individual proteins at pH 6.0 calculated using SnapGene program along with the number of hydrophobic residues and hydropathy indices estimated using the CIDER platform.

|  | *Charge*<br>*(pH 6.0)* | *Net charge per residue*<br>*(pH 6.0)* | *Hydrophobic*<br>*residues* | *Hydropathy*<br>*index* |
| --- | --- | --- | --- | --- |
| GRN-2 | -4.21 | -0.07 | 23 | 4.68 |
| GRN-3 | -2.03 | -0.03 | 21 | 4.40 |
| GRN-5 | -6.00 | -0.10 | 17 | 4.38 |
| GRN-7 | +2.78 | +0.04 | 16 | 3.90 |
| PrLD | +7.63 | +0.04 | 27 | 3.81 |

## Notes

### Competing Interest Statement

The authors have declared no competing interest.

